# Co-infection with *Toxoplasma gondii* leads to a loss of resistance in *Heligmosomoides bakeri* trickle-infected mice due to ineffective granulomas

**DOI:** 10.1101/2020.12.17.423220

**Authors:** Breton Fougere, Anupama Ariyaratne, Naomi Chege, Shashini Perera, Emma Forrester, Mayara de Cassia Luzzi, Joel Bowron, Aralia Leon Coria, Edina Szabo, Constance A. M. Finney

## Abstract

The intestinal roundworm *Heligmosomoides bakeri* causes chronic infection in susceptible (C57Bl/6) mice; however, repeat (trickle) infection confers immunity and facilitates worm clearance. We previously showed that this acquired immunity is associated with a strong Th2 response, notably the enhanced production of intestinal granulomas. Here we demonstrate that elevated proportions of IgG_1_-bound eosinophils and macrophages are observed around the developing tissue worms of trickle-infected female C57Bl/6 mice compared to bolus infected animals. Levels of IgG_2c_, IgA or IgE were not detected in the granulomas. Increased proportions of SiglecF^+^ and CD206^+^ cells, but not Ly6G^+^ and/or NK1.1^+^ cells, were also found in the granulomas of trickle-infected mice. However, in the natural world rather than the laboratory setting, immune environments are more nuanced. We examined the impact of a mixed immune environment on trickle infection-induced immunity, using a pre-infection with *Toxoplasma gondii*. The mixed immune environment resulted in fewer and smaller granulomas with a lack of IgG_1_-bound cells as well as reduced proportions of SiglecF^+^ and CD206^+^ cells, measured by immunofluorescence and flow cytometry. This was associated with a higher worm burden in the co-infected animals. Our data confirm the importance of intestinal granulomas and parasite-specific antibody for parasite clearance. They highlight why it may be more difficult to clear worms in the field than in the laboratory.

**AUTHOR’S SUMMARY:** Despite decades of research on intestinal parasitic worms, we are still unable to clearly point to why so many people (approximately 1.8 billion) and most livestock/wild animals are infected with these parasites. We have made progress in understanding how the immune system responds to parasitic worms, and how these parasites manipulate our immune system. However, identifying effective clearance mechanisms is complex and context dependent. We have used models of trickle infection (multiple low doses of parasites) and co-infection (two intestinal parasites) to simulate how people/animals get infected in the real world. Using these models, we have confirmed the host/parasite interface (the granuloma) within the intestinal tissue to be key in determining the host’s ability to clear worms. The lack of specific immune cells and antibodies within the granuloma was associated with chronic infection. Our results help explain why intestinal parasitic worms are so prevalent and why it may be difficult to clear worms in natural settings.

## INTRODUCTION

Gastrointestinal nematodes (GINs) are ubiquitous. They are highly prevalent in humans ^1^, livestock (reviewed in ^2^) and pets ^3^. Despite the low mortality associated with these parasites, they significantly impact human and animal health (reviewed in ^4–6)^. Infected hosts mount strong anti-GIN immune responses, but the efficacy of the response is dependent on genetics, infection dynamics and the environment, and we have yet to find treatments that clear worms and prevent reinfection. Understanding effective clearance mechanisms by the host in natural settings is key to developing strategies to stimulate sterile immunity.

*Heligmosomoides bakeri* (Hb), formerly *Heligmosomoides polygyrus* ^7^, is a rodent model gastrointestinal nematode parasite which has been extensively used to understand host/parasite interactions in related livestock and human parasites ^8^. For the first 7-10 days of infection, Hb develops within the tissue. After 10 days, adults emerge into the lumen, where they remain coiled around the intestinal villi for the duration of infection. Hb infection stimulates a strong Th2 response, ultimately dependant on the production of IL-4 and IL-13, and IL-4 receptor signalling for effective clearance ^9,10^. Tissue dwelling parasite stages induce an influx of immune cells to the site of infection culminating in the formation of an intestinal granuloma (reviewed in ^11^). The levels of IL-4/IL-13, and the size and number of granulomas are associated with the resistant/susceptibility phenotype of different mouse strains ^12,13^.

Immune protection from reinfection, observed in laboratory models of vaccination and/or secondary infections, has been linked to effective immune responses against tissue dwelling Hb stages ^14–16^. Within the granuloma, the host response simultaneously immobilises, damages and/or kills the parasites while healing the damage caused by the growing worm (reviewed in ^11^). Due to their size, parasitic worms cannot be phagocytosed by immune cells; antibody-bound myeloid cells, namely SiglecF+ eosinophils and CD206+ macrophages, within the granuloma have been implicated in protection (reviewed in ^11^).

Resistant strains of mice develop faster and stronger parasite-specific antibody responses to Hb, as compared to susceptible strains ^12,13^. More specifically, elevated levels of IgG, IgA and IgE have been linked to worm clearance ^17–20^. Passive transfer of serum, more specifically IgG_1_, from infected mice decreases adult worm numbers as well as their fitness ^21–24^. Mice lacking IgG, and to a lesser extent IgA, but not IgE, have increased worm burdens upon secondary infection compared to wild-type animals ^18^. In support of this, macrophages isolated from Hb-induced granulomas from challenge infected mice are bound to IgG isotypes (IgG and IgG) but not IgE ^15^. They also express high levels of FcγRs, and mice lacking FcγRs (unable to bind IgG antibodies) have increased parasite loads ^16^.

While it’s clear that in the context of a Th2 immune environment generated in a laboratory-based experimental model, susceptible strains of mice can mount strong immune responses to reinfections with Hb ^13,18^, in natural settings, where immune environments are more nuanced, Hb prevalence is high ^25–27^. One of the many factors that influence immune environments is co-infection, a common occurrence in the natural world ^28^.

The overarching aim of our study was to examine the impact a mixed immune response on effective clearance of Hb. As an obligate intracellular parasite, *Toxoplasma gondii* (Tg) elicits a strong inflammatory response in the small intestine during infection, with elevated IL-12 and IFNγ levels (as reviewed in ^29^). Following infection of the intestinal mucosa, IL-12 producing monocytes and neutrophils are recruited to the small intestine, where they recruit ILC1s and T cells, both potent producers of IFNγ. Research suggests that previous Tg infection skews Hb-specific CD4^+^ T cell and IgG antibody responses towards a more Th1 phenotype, with no impact on worm burden ^30^. However, this previous study focused on a primary chronic infection (28 days post infection - D28) and did not address potential impacts to the host/parasite interface in the granuloma. We used the more realistic trickle model of TgHb confection and show that TgHb infected animals have a reduction in the size and number of granulomas, and that their granulomas have fewer myeloid cells and more specifically, IgG_1_-bound myeloid cells. The lack of functional granulomas was associated with increased worm burdens. Our data confirm the importance of intestinal granulomas in the context of re-infection, and highlight why it may be more difficult to clear worms in the field than in the laboratory.

## MATERIALS AND METHODS

### Mice, Parasites and Antigens

Female BALB/c and C57Bl/6 mice aged 6-10 weeks were bred and maintained in the Life and Environmental Science Animal Resource Centre (LESARC) in the Department of Biological Sciences, University of Calgary, Canada. Original stocks were obtained from Charles River, Canada. Mice were acclimatised for a minimum of two days when moved to our experimental room and housed in groups of 3-5 in individually vented cages. All experiments were approved under protocol # AC17-0083 and AC19-0121 by the University of Calgary’s Life and Environmental Sciences Animal Care Committee. All protocols for animal use and euthanasia were in accordance with the Canadian Council for Animal Care (Canada). Enrichment was provided in the form of plastic housing, bedding and for parasite passage animals, a variety of veterinary approved small treats (seeds, nuts, etc) and/or supportive nutrition (DietGel).

Mice were orally infected by gavage with 200 third stage *Heligmosomoides bakeri* larvae (maintained in house, stock was a gift from Dr. Allen Shostak, University of Alberta, Canada and Dr Lisa Reynolds, University of Victoria, Canada) and euthanized at either D5, D7, D21 or D28. Larvae were obtained from fecal cultures after approximately 8-9 days of incubation at room temperature. Mice were infected according to the bolus or trickle infection regimes (Figures 1A & 4A). To avoid differences in counts during the trickle infections, on D0, two identical solutions were made up (200 worms/100μl). One was used to infect the bolus infected mice on D0 and one was used for the trickle-infected mice. The solution for the trickle-infected animals was divided into ten equal parts. Each part was made up to 100ul using water. Animals were gavaged with a diluted solution on days 0, 2, 4, 6, 8, 10, 12, 14, 16 and 18. Using the 10 doses, trickle-infected animals received 200 larvae in total over 18 days.

**Figure 1:**
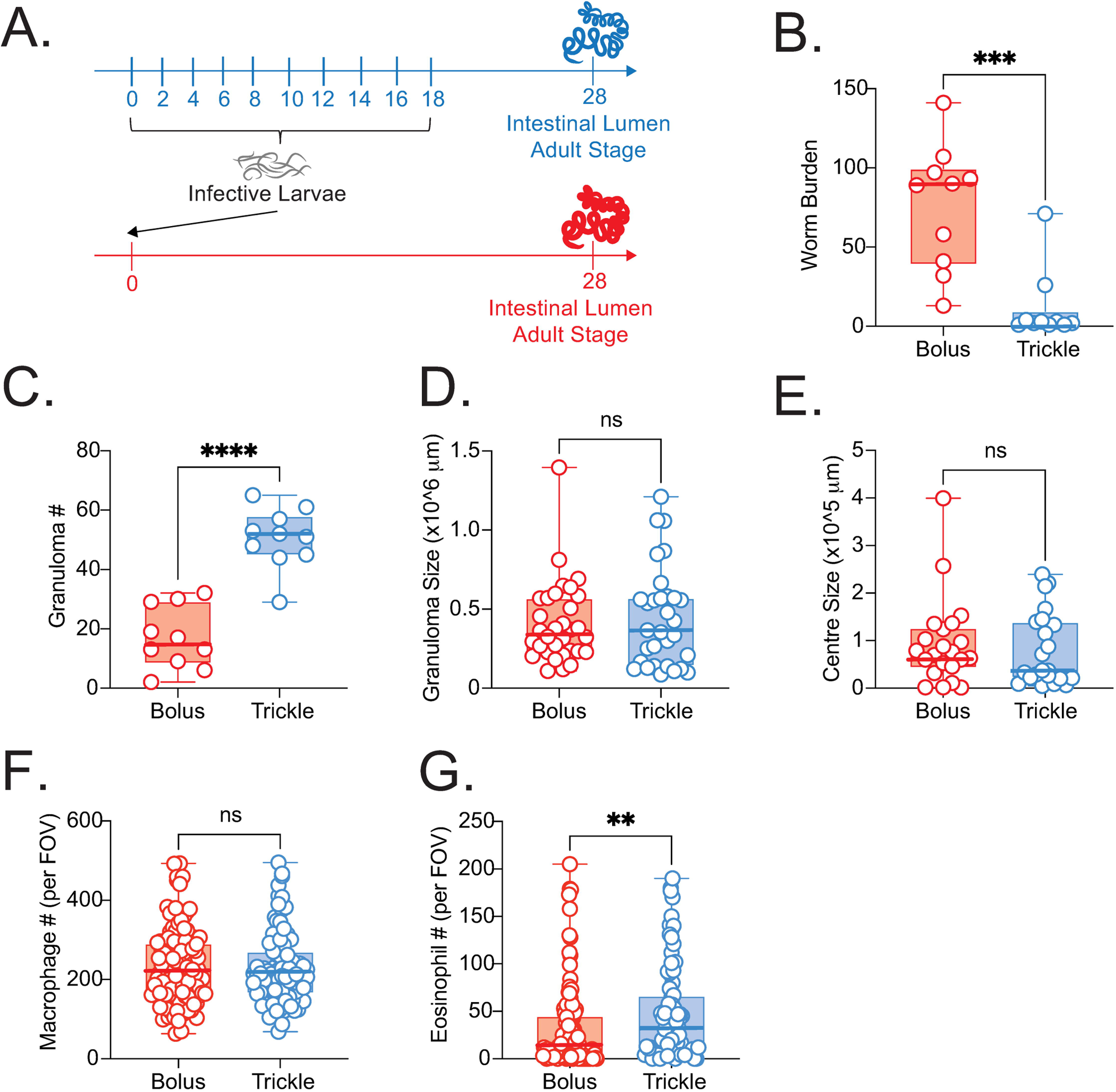
Characteristics of the H. bakeri trickle infection model in C57Bl/6 mice. C57Bl/6 mice were infected with 200 *H. bakeri* according to the bolus and trickle infection regimes. (A) Trickle infection regime: mice are infected with 200 larvae in total, but in multiple doses over the course of infection (in blue). Doses of 20 larvae are trickled on days 0, 2, 4, 6, 8, 10, 12, 14, 16 and 18 post-infection This leaves a 10-day window after the final dose to allow parasites to fully develop into adults and migrate from the intestinal tissue to the intestinal lumen. Bolus infected mice were infected with 200 larvae at day 0 (in red). Adult worms (B) and granulomas (C) were counted in the small intestine at D28. Mice were infected according to the bolus (red) and trickle (blue) regimes. Total granuloma area (D) and the granuloma necrotic centre (E) were measured on H and E stained slides of whole small intestine. Macrophages (F) and eosinophil (G) were counted within the center of the granulomas at x400 magnification. Cells were counted for one field of view (FOV) per granuloma. Graphs represent pooled data from a minimum of 2 experiments, bars represent the median, with a minimum of 3 mice per group per experiment. A normality test (D’Agostino & Pearson) was performed, followed by a Kruskal Wallis/Anova (with post-tests) and Mann Whitney/T-test to test for statistical significance between trickle and bolus groups, n.s. = not significant, *** = p<0.001, **** = p<0.001.

The Me49 strain of *Toxoplasma gondii* (Tg) (maintained in house, original stock was a gift from Dr. Georgia Perona Wright, University of British Columbia, Canada) was maintained in male C57BL/6 and CD1 mice by alternating bimonthly passage into new animals. Cysts were obtained from the brains of these animals a month after infection. Ten cysts were orally administered per mouse (female C57BL/6) followed by Hb larvae 14 days later. Mice were euthanized 21 or 28 days post Hb infection (Fig 7A).

Sample size was calculated based on numbers used in our previous work ^13^ or for the flow cytometry of granuloma cells, on infecting enough animals to obtain the required amount of cells. No animals for which we obtained data were excluded from analysis. However, criteria had been set. We would have excluded all animals developing pathologies unrelated to infections during the experiment or observed during necropsy (e.g. tumours). Animals were not randomised. Co-infections and trickle infections require many manipulations. Our experience is that using confounders increases mistakenly providing the incorrect treatment to the mice.

All mice were monitored daily. Humane endpoints were defined as: general decline of health, respiratory distress, severe neurological signs, loss of body weight > 20% or recommendation from the facility veterinarian. Animals were euthanised within 24 hours of humane endpoints being observed. For TgHb animals, a small number of animals were found dead before the 28 days post-Hb infection. This happened towards the end of the experiment, usually in the early morning, after a check of humane endpoints the previous day.

Hb antigen was prepared by collecting live adult worms from 14/21 or 28-day infected mice using modified Baerman’s apparatus. Worms were washed multiple times and homogenized in PBS using a glass homogenizer. The resulting solution was centrifuged (13, 000 g, 10 minutes, 4°C) and the supernatant filtered (0.2mm filter, Nalgene). The protein concentration was calculated using the Bradford assay. The antigen was stored at 15 mg/ml at −80°C.

We collected immune serum from trickle infected mice euthanized at D14 using a previously published trickle infection model ^13^. We transferred 100ul intravenously to bolus infected mice at day 0, day 2, day 4 and day 6.

### Worm and Granuloma Counts

Small intestines of infected mice were harvested and opened longitudinally. The number of adult worms present in the intestinal lumen along the length of the small intestine was counted under a dissection microscope. Macroscopic granulomas were counted by eye under a dissection microscope with the small intestine intact.

### Granuloma size

Consecutive formalin fixed paraffin embedded mouse small intestinal sections were stained with H & E by the University of Calgary’s Veterinary Diagnostic Services Unit. Stained tissue sections were analyzed visually using the Leica DMRB light microscope and/or the OLMYPUS SZX10 microscope. Images of granuloma were captured using the QCapture and the cellSens software and processed using the ImageJ software, in a blinded manner. Whole granuloma and necrotic centre size were measured for each granuloma on the slide. Granuloma area was calculated using ImageJ. Granuloma were outlined using the freehand perimeter tool and area was calculated using a pre-determined measurement scale.

### Serum ELISAs

Blood samples were left to clot for 30 minutes and then centrifuged twice at 11, 000 g at 4°C for 10 minutes. Serum was collected and used either fresh or stored at −80°C. Total IgE (BD, 555248) and IgA (capture antibody, BD, 556969, detection antibody, BD, 556978) and IgG1 levels (capture antibody, BD, 553440, detection antibody, BD, 553441) were measured by ELISA according to manufacturer’s instructions.

Antigen specific antibody responses were also measured by ELISA. ELISA microplates were coated with 10 mg/ml *H. bakeri* antigen in carbonate buffer (0.1mM NaHCO3, pH 9.6), overnight at 4°C. Plates were blocked with 2% BSA in TBS/ 0.05% tween 20 for 2 hours at 37°C. Sera were diluted in TBS Tween and added to wells overnight at 4°C. Antigen-specific IgG1 was detected with HRP-conjugated corresponding detection antibodies (anti-IgG1, BD, 553441) with TMB peroxidase substrate (T3405, Sigma). The reaction was stopped using 1M H2SO4 solution and the color change was read at 450nm.

### Intestinal tissue homogenate ELISAs

Small intestines were opened longitudinally and washed with PBS to remove luminal content. The mucosal surface was identified under a dissecting microscope. The mucosal surface (with its mucus) was gently scraped using a glass slide. Scrapings were weighed, added to 500 ml lysis buffer (10 mM tris HCl, 0.025% sodium azide, 1% tween 80, 0.02% phenylmethylsulfonyl fluoride) with one complete protease inhibitor tablet (Roche diagnostics GmbH, Germany) and homogenized using a bead beater (40 seconds at speed 6 using the Fast-prep-24 bead beater, MP biomedical). The homogenate was centrifuged at 11, 000 g at 4°C for one hour. Supernatants were collected and used fresh or stored at −80°C. IgA (IgA (capture antibody, BD 556969, detection antibody, BD 556978), IL-4 (R & D systems, DY404 kit), IL-13 (R & D systems, DY413 kit) and IL-21 (R & D systems, DY594 kit) was measured by ELISA according to manufacturer’s guidelines. All IL-21 ELISAs were performed on fresh, unfrozen material ^31^.

### MLN and SPL Cell isolation, *in vitro* re-stimulation assay and cytokine ELISAs

Mesenteric lymph nodes (MLN) and spleens (SPL) were mechanically dissociated into single cell suspensions. Cells were counted using a Beckman-Coulter ViCell XR. MLN and SPL were cultured at 1 × 10^6^ cells/ml for 48 hours in RPMI medium, 10% FCS, 1% L-glutamine, 1% penicillin/ streptomycin (supplemented RPMI 1640) in the presence of 10 mg/ml *H. bakeri* antigen or 2 mg/ml concanavalin A (Sigma) at 37°C with 5% CO2. Supernatants were collected for cytokine measurements. Measurements for antigen-specific production were not included in the analysis unless cytokine production was observed in the wells with concanavalin A stimulation. ELISAs were performed according to manufacturer’s guidelines (R & D systems, DY404 kit for IL-4 and DY413 kit for IL-13).

### Transit time

Gastrointestinal transit time was measured one day prior to euthanasia. Mice were fasted for 6 hours and 200 ml of 5% Evans blue (Sigma) in 5% gum arabic (ACROS organics) was orally gavaged using a ball tip 20 gauge 1.5’’, 2.25mm curved animal feeding needle. Each mouse was labelled, with the time of dye administration recorded. Mice were transferred to clean empty cages and the time to pass the first blue fecal pellet was recorded. Gastrointestinal transit time was calculated for each mouse.

### Immunofluorescence

Whole small intestines were fixed in formalin for a minimum of 48 hours, washed in saline for a minimum of 48 hrs and processed to obtain paraffin embedded tissue blocks. 6um sections were cut by the Diagnostic Services Unit, UCalgary. The sections were stained for H&E or with either anti-mouse IgG_1_ (BD 562026), anti-mouse IgA (BD 559354), anti-mouse IgE (BD 553415), anti-mouse IgG_2c_ (Invitrogen SA5-10221) or their respective isotypes overnight at 4°C. Slides were mounted with FluoroshieldTM with DAPI (Sigma-Aldrich F6057). Images were acquired using the Zeiss Axio ZoomV.16 stereoscopic microscope (Bio Core Facility, University of Calgary) or the Thorlabs Tide whole-slide scanning microscope (Live cell imaging Facility, University of Calgary). Brightfield and fluorescence images were taken either under the 2.3x objective with the AxioCam High Resolution color (HRc) and High-Resolution mono (HRm) cameras, or the 20x/0.75 NA objective of the slidescanner respectively. Zeiss Zen (blue edition) software or Thorlabs Tide LS image acquisition software was used. DAPI, FITC and Texas Red fluorophores were used. Brightness and contrast were adjusted similarly for all photographs in photoshop.

### Granuloma Digestion for Flow cytometry

Granulomas were identified along the small intestine as tissue (containing worms) or lumen (chronic granuloma not containing worms) using a dissection microscope. Granulomas from 5 mice were pooled by infection type (trickle/bolus/co-infected) and granuloma type (tissue/lumen). Isolated granulomas were incubated with DNAse (Roche 10104159001) and 1:50 Liberase (Roche 0540102001)) at 37°C for 90 minutes. After incubation, granulomas filtered, washed, centrifuged and stored at 4°C until use. Cells were counted using a Vi-CELLTM XR Cell Analyzer (Beckman Coulter 731196). 1×10^6^ cells (100μL) were used per sample. Samples were stained with fixable viability stain (BD, 565388 or BD 564406) for 15 minutes at RT in dark, blocked with rat anti-mouse CD16/32 (BD 553142) for 5 minutes at 4°C in the dark and stained for surface markers for 20 minutes at 4°C in the dark. For the granuloma panel, cells were stained for CD3 (BD 560591), CD19 (BD 612781), NK1.1 (BD 564143), SiglecF (BD 552126), CD206 (BD 565250), Ly6G (BD 740953), CD11b (BD 563553), and IgG1 (BD 553443).

For intracellular staining (IgG_1_ and IgA), samples were fixed, permeabilized and stained for IgG_1_ (BD 553443) and IgA (BD 743293) or isotype controls (BD 554684, BD 562868) for 30 minutes at 4°C. Samples were then washed and centrifuged. Resuspended samples were run on an LSRFortessaTM X20 (BD Biosciences) flow cytometer. Data was acquired using FACSDivaTM software, and analysis of cell populations was performed using FlowJoTM software (v10.7.1).

### MLN and Peyer’s Patches Isolation for Flow Cytometry

Peyer’s patches (PP) and MLN were removed from the small intestine under the dissection microscope. Cells were mechanically dissociated into single cell suspensions. PP and MLN cells were counted using a Vi-CELLTM XR Cell Analyzer (Beckman Coulter 731196). 1×10^6^ cells (100μL) were used per sample. Samples were stained with fixable viability stain (BD 564406) for 15 minutes at RT in dark, blocked with rat anti-mouse CD16/32 (BD 553142) for 5 minutes at 4°C in the dark and stained for surface markers for 20 minutes at 4°C in the dark. For the plasma cell panel, cells were stained for CD3 (BD 566495), IgD (BD 558597), B220 (BD 561880), CD19 (BD 612781), and CD138 (BD 563193/565176).

### RNA/DNA Extraction and Quantitative PCR

RNA and gDNA were extracted from 4 equal snap frozen sections of the SI (sections 1-4, where 1 is the most proximal section and 4 the most distal from the stomach) by crushing the tissue in Trizol (Ambion, Life Technologies) using a pestle and mortar on dry ice. We followed manufacturer guidelines for RNA extraction. To measure cytokine levels in the RNA samples, cDNA was prepared using DNase 1 (RNase-free, New England Biolabs), and iScript Reverse Transcription Supermix for RT-qPCR (Bio-Rad) or AdvanTech 5x Revese Transcription master mix (Diamed).

Primers for quantitative PCR were obtained from Integrated DNA Technologies (San Diego) for IL-13 (GATCTGTGTCTCTCCCTCTGA forward, GTCCACACTCCATACCATGC reverse), IFNγ (TCAAGTGGCATAGATGTGGAAGAA forward, TGGCTCTGCAGGATTTTCATG reverse), IL-10 (GTCATCGATTTCTCCCCTGTG forward, ATGGCCTTGTAGACACCTTG reverse) and 18S rRNA (GCAATTATTCCCCATGAACG forward, GGCCTCACTAAACCATCCAA reverse). Samples were run in triplicate on a Bio-Rad CFX96 real time system C1000 touch thermocycler. Relative quantification of the cytokine genes of interest was measured with the delta-delta cycle threshold quantification method ^32^, with 18S rRNA for normalization. Data are expressed as a fold change relative to 18S rRNA levels for each animal.

### ELIspot Assay

Plates were then coated with either purified rat-anti mouse IgG1 (BD 553440) or Hb antigen and incubated overnight at 4°C. Plates were washed and blocked with media for 2 hours at 37°C with 5% CO_2_. MLN cells added in triplicate at 5×10^3^ cells/mL to measure total IgG1 producing cells. For quantifying *H. bakeri* antigen-specific IgG1 producing cells, a total of 2×10^6^ cells were cultured overnight at 37°C with 5% CO_2_. For detection, IgG1 (BD 553441) was incubated for 2 hours at RT, followed by a Streptavidin-HRP (horseradish peroxidase; BD 554066) incubation for 1 hour at RT. Spots were allowed to develop for 5-7 minutes once the substrate solution had been added, before stopping the reaction. Plates were allowed to dry for 24-48 hours, before spots were counted manually under a dissecting microscope.

### Statistical Analysis

Graphpad Prism software (La Jolla, CA, USA) was used for all statistical analysis. To compare multiple groups (three or more), we performed a normality test (D’Agostino and Pearson test, unless N too small, in which case Anderson-Darling or Shapiro-Wilk). ANOVA or Kruskal-Wallis tests were performed on parametric/non-parametric pooled data, and when significant, Sidak’s/Dunn’s Multiple comparisons were performed on Hb vs. HbTg and Tg vs. HbTg. For comparisons of 2 groups, statistical significance was assessed by either an unpaired t test (for data sets with normally distributed data) or Mann-Whitney’s U test (for non-parametric data) or paired t test (for paired data sets with normally distributed data). Data are presented as median and individual data points, unless otherwise specified.

## RESULTS

### The resistant phenotype in trickle infected C57Bl/6 mice is not associated with increased Th2 cytokine or antibody levels

We have previously shown that using a trickle regimen, C57Bl/6 mice harbour fewer Hb worms and have more granulomas than bolus infected mice ^13^. Using a more robust trickle model (more doses and later time points, Fig. 1A), we show even starker differences depending on the mode of infection. After 28 days (D28), the Hb worm burden (primary outcome) of the trickle-infected group was below 5 worms in 8/10 animals compared to a mean of 76 in bolus infected animals (Fig 1B). The granuloma number was also higher in trickle infected animals, with 9/10 mice having more granulomas than the bolus infected mouse with the most granulomas (32 granulomas, Fig. 1C). Despite granuloma size being associated with susceptibility/resistance phenotypes ^12^, we found that this parameter was not affected by the mode of infection. Neither the size of the whole granuloma, nor the size of the internal necrotic centre changed with the mode of infection (Fig 1D and 1E). However, as we have previously shown with trickle infection ^13^, granuloma composition differs between bolus and trickle infected animals. Macrophages numbers are similar (Fig 1F) but eosinophil numbers are increased (Fig 1G). In resistant Balb/c animals, we see no differences in worm burden or granuloma number between bolus and trickle infected animals (Sup Fig 1A and 1B).

A strong Th2 response is responsible for anti-parasitic worm immunity. Increased levels of IL-4 and IL-13 in the MLN (mesenteric lymph nodes) and SPL (spleen) are commonly reported in bolus infected animals and have been directly linked to parasite clearance^9,10,33–35^. However, in trickle infected animals, previous studies have not linked reduced parasite burdens to an increase in these cytokines ^13,36^. We found no differences in the levels of IL-4 and/or IL-13 between bolus and trickle infected mice, apart from IL-13 in the SPL (Fig 2A-F). SPL IL-13 levels were recorded per cell and likely decreased in the trickle infected animals in response to the low worm burden. When accounting for differences in total cell number (Fig 2E and 2F), IL-4 and IL-13 levels in the MLN and SPL in trickle and bolus infected animals remain similar. Levels of IL-4 and IL-13 were undetectable in the serum, as previously observed ^13^. Again, in resistant Balb/c animals, we see no differences between bolus and trickle infected animals in any of the parameters measured, apart from spleen IL-13 (Sup Fig 1C-H).

**Figure 2:**
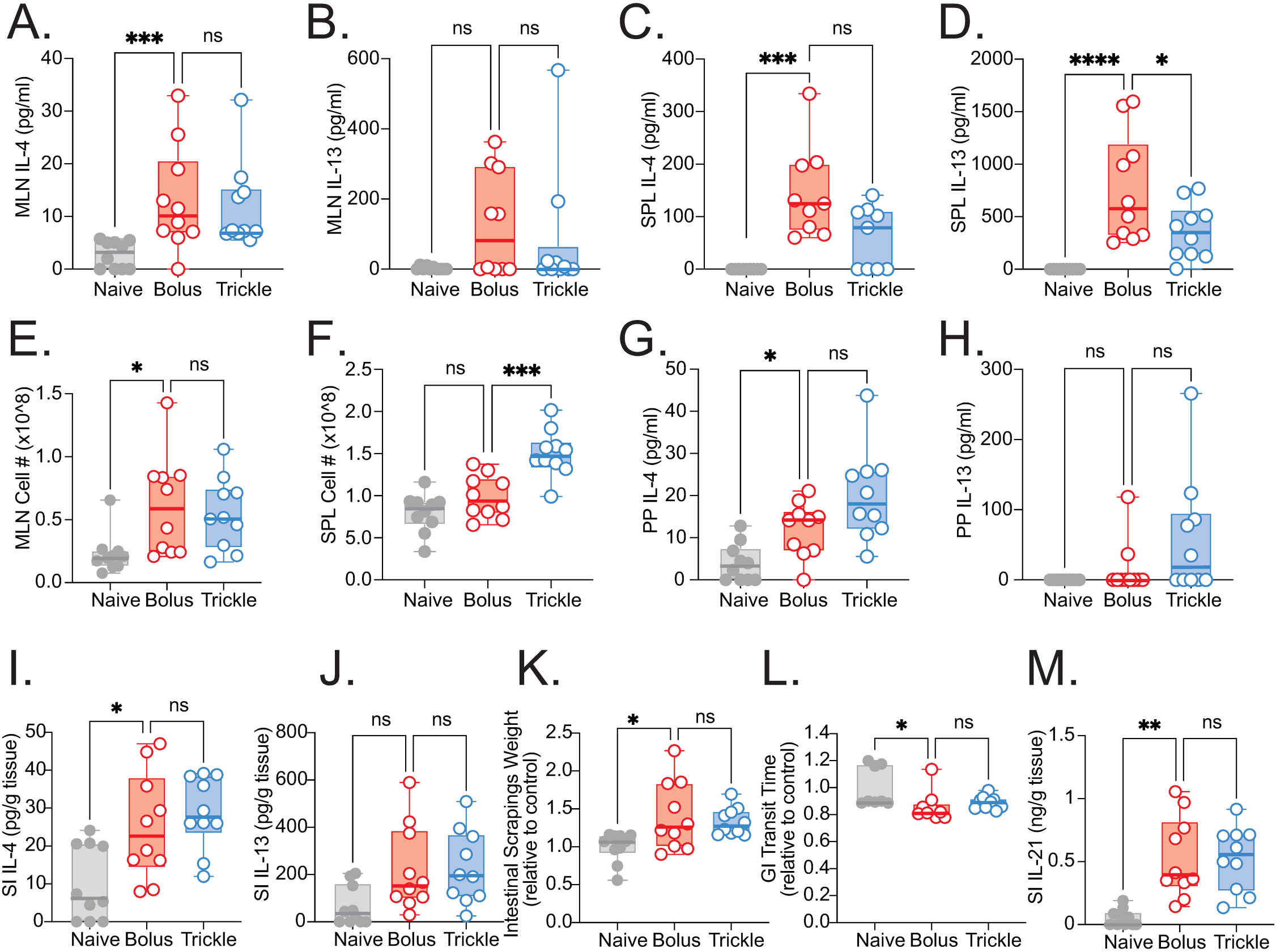
Th2 response in the SPL, MLN, PP and SI during H. bakeri trickle infection in C57Bl/6 mice. C57Bl/6 mice were infected with 200 *H. bakeri* larvae according to the bolus and trickle infection regimes. At D28, single cell suspensions were isolated from the spleen (SPL), mesenteric lymph nodes (MLN) and Peyers Patches (PP) and cultured for 48 hours in the presence of Hb antigen. (A-D) IL-4 and IL-13 MLN and SPL cytokine levels were measured in the supernatant by ELISA. Total cell numbers were calculated for the MLN (E) and SPL (F). IL-4 (G) and IL-13 (H) PP cytokine levels were measured in the supernatant by ELISA. (I-M) Small intestines were dissected and scraped using a glass slide leaving only the serosa. (I-J) IL-4, and IL-13, were measured in the intestinal scrapings by ELISA. (K) The intestinal scrapings were weighed for each mouse and normalised to control animals. (L) Mice were fasted for 6 hours followed by Evans Blue administration. Time from dye administration to the passing of dyed fecal pellets was measured and normalised to control animals. (M) IL-21 was measured in the intestinal scrapings by ELISA. Mice were infected according to the bolus (red) and trickle (blue) regimes. Naive mice are represented by grey circles. Graphs represent pooled data from 2 experiments, bars represent the median, with a minimum of 3 mice per group per experiment. A normality test (D’Agostino & Pearson) was performed, followed by a Kruskal Wallis/Anova (with post-tests) for statistical significance between trickle and bolus groups, n.s. = not significant, * = p<0.05, ** = p<0.01, *** = p<0.001, **** = p<0.001.

Th2 immune responses can also be observed closer to the host/parasite interface, in the Peyer’s patches (PP) and the small intestine (SI). Levels of the Hb antigen-specific IL-4 and IL-13 were again similar between bolus and trickle infected animals (Fig 2G-J). No levels of IL-5, IL-9 or IL-10 were detectable in intestinal tissue from control or infected mice, as previously observed ^13^. Increased mucus production ^37^, intestinal smooth muscle contractility ^38^ and IL-21 levels ^39^ have all been associated with parasite clearance. However, none of these parameters differed between the two infected groups in the SI (Fig. 2K-M).

Resistance to Hb has also been linked to increased parasite-specific IgG_1_, IgA and IgE levels ^18^. Total IgG_1_ and IgE levels were increased in the serum of infected vs. naïve mice (Fig 3A and 3B). However, the levels did not change with the mode of infection. No difference was observed in the levels of parasite specific IgG_1_ either (Fig 3C). Levels of intestinal IgA did not differ between naïve, bolus infected or trickle infected animals (Fig 3D). No detectable levels of Hb larval or adult parasite antigen specific IgE, IgG_2c_ or IgA were observed in the serum of bolus-or trickle-infected mice. In resistant Balb/c animals, we see no differences between bolus and trickle infected animals in IgE or IgG_1_ levels (Sup Fig 1I-K).

**Figure 3:**
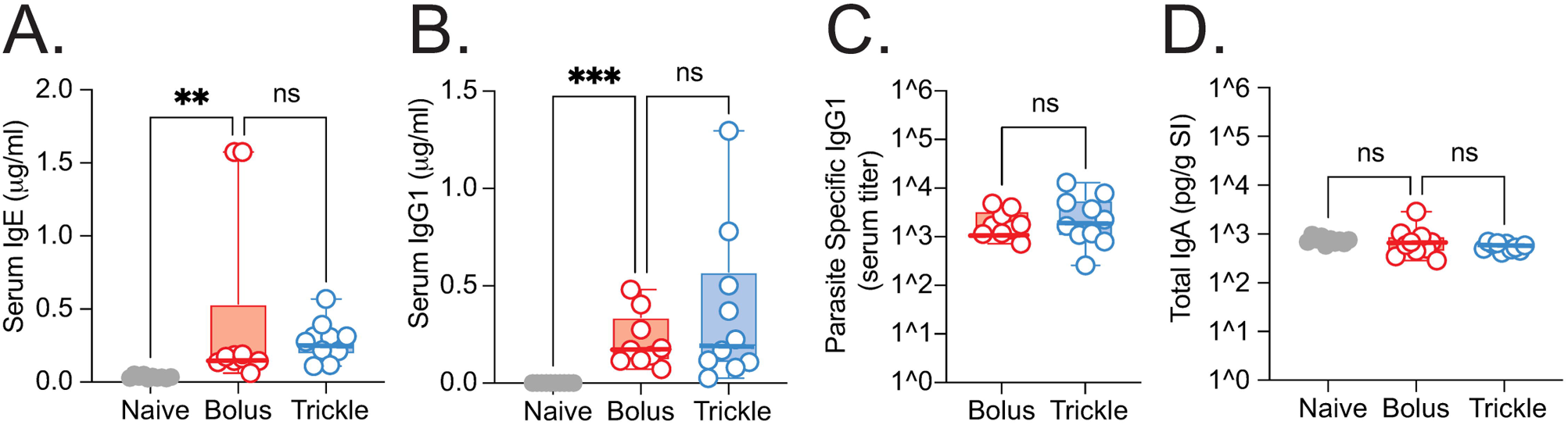
Antibody levels during H. bakeri trickle infection in C57Bl/6 mice. C57Bl/6 mice were infected with 200 *H. bakeri* larvae according to the bolus and trickle infection regimes. At D28, blood was collected and serum obtained. (A) Total serum IgE, (B) Total serum IgG_1_, (C) Parasite-specific serum IgG_1_ and (D) Total intestinal IgA were measured by ELISA. Levels in naïve controls were undetectable for parasite specific IgG_1_. Graphs represent pooled data from a minimum of 2 experiments, bars represent the median, with a minimum of 3 mice per group per experiment. A normality test (D’Agostino & Pearson) was performed, followed by a Kruskal Wallis/Anova (with post-tests) and Mann Whitney/T-test to test for statistical significance between trickle and bolus groups, n.s. = not significant, ** = p<0.01, *** = p<0.001.

By D28 of infection, all worms had migrated from the tissue to the lumen. Our results are therefore unsurprising in light of the importance of immunity against developing tissue parasite stages, which are best observed at the host/parasite interface early during infection. We next focused our efforts on studying immune responses within the granuloma during acute infection.

### Cell-bound IgG_1_ antibodies accumulate around encysted larvae in the granulomas of trickle-infected C57Bl/6 mice

By D10 most, if not all, parasites have matured into adults, and left the tissue to inhabit the lumen. Previous research has demonstrated that the resistance to Hb attained by the host after vaccination and/or secondary infection is aimed at incoming larvae ^14,15^. To study immune responses against larvae in the trickle infected animals, we euthanised mice at D21 instead of D28 (3 instead of 10 days after the final parasite dose). This ensured there were both developing parasites in the tissue and adults in the lumen. We matched these to an early (D5) and late (D21) bolus infection (Fig 4A).

**Figure 4:**
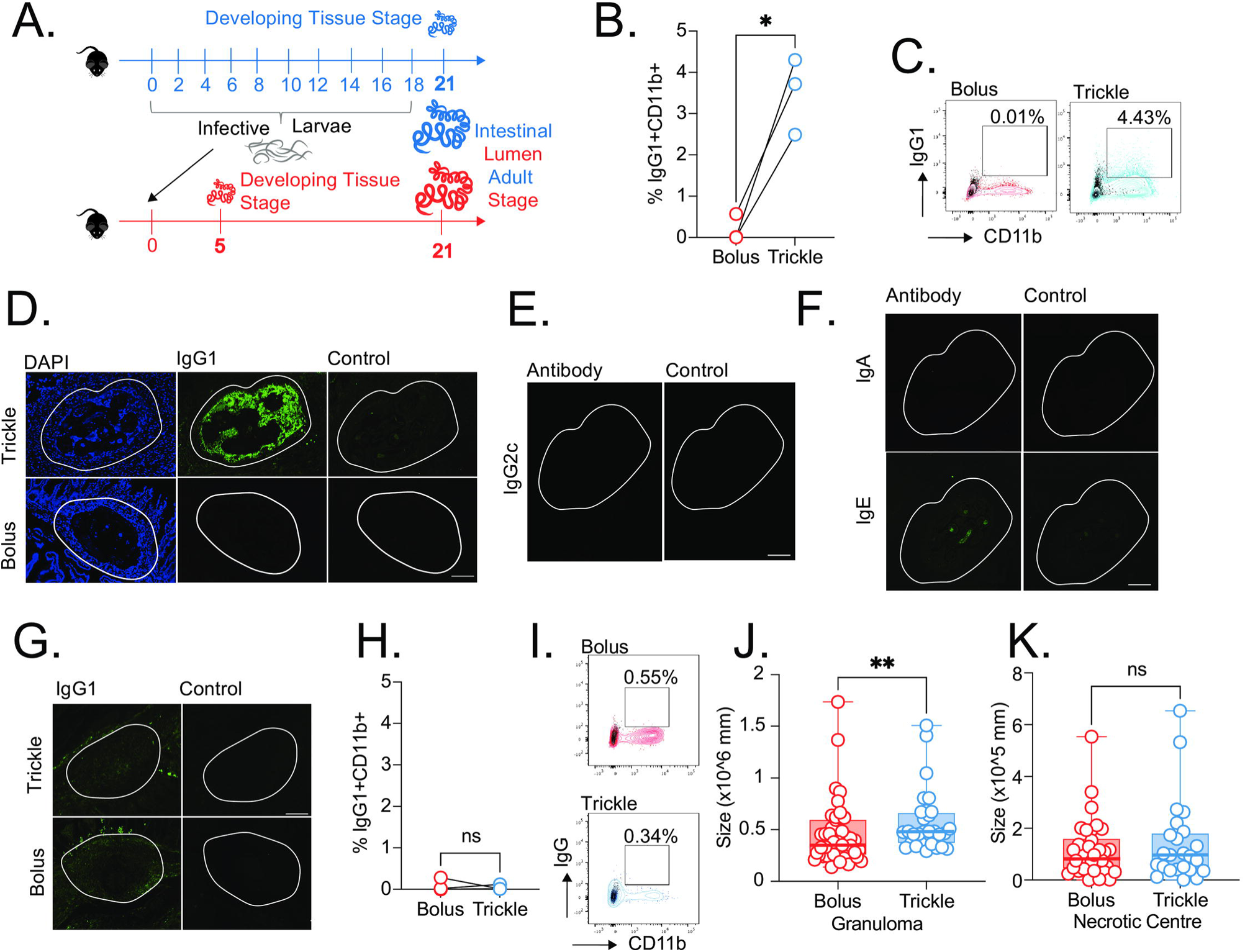
IgG_1_ in the granulomas of trickle-H. bakeri infected C57Bl/6 mice. C57Bl/6 mice were infected with 200 *H. bakeri* according to bolus and/or trickle models of infection. (A) Trickle infection regimen (worm-containing granulomas): mice are infected with 200 larvae in total but spread out over infection (in blue). Mice are euthanised on day 21, doses of 20 larvae are trickled on days 0, 2, 4, 6, 8, 10, 12, 14, 16 and 18 post-infection. There is a 3 day window after the final dose to allow parasites to start developing in the mucosa and observe the formation of granulomas around them, as well as having mature worms in the lumen. For the comparative bolus infections, we are using animals infected for 5 (tissue worms only) and 21 (worms found in the lumen only) days. (B & H) Granulomas were isolated from the small intestine of 5 mice per group. Granulomas from the individual animals were pooled to form one sample per group. Cells were obtained by tissue digestion of the pooled samples and stained by flow cytometry. (B) The percentage of CD11b^+^IgG ^+^ cells as a proportion of live granuloma cells (with T and B cells removed) from acute granulomas was calculated for three independent experiments. (C) Representative flow cytometry plots showing IgG_1_ and CD11b expression on live granuloma cells from acute granulomas. (D-G) Formalin fixed, paraffin-embedded sections were obtained from small intestine swiss rolls. These sections were co-stained with DAPI and either anti-mouse (D) IgG_1_, (E) IgG_2c_, (F) IgA/IgE and (G) IgG_1_. Scale bar=100μm. Antibody stain (left), isotype control stain (right). Representative granulomas (white lines) containing worms (acute granulomas, D-F) or not containing worms (chronic granulomas, G) from trickle and bolus infected mice. (H) The percentage of CD11b^+^IgG ^+^ cells as a proportion of live granuloma cells (with T and B cells removed) from chronic granulomas was calculated for three independent experiments. (I) Representative flow cytometry plots showing IgG_1_ and CD11b expression on live granuloma cells from chronic granulomas. (J) Total granuloma area (K) and the granuloma necrotic centre were measured for chronic granulomas on H and E stained formalin fixed sections from whole small intestine swiss rolls. Graphs represent pooled data from a minimum of 2 experiments, bars represent the median, with a minimum of 3 mice per group per experiment. A normality test (D’Agostino & Pearson) was performed, followed by Mann Whitney/T-test to test for statistical significance between trickle and bolus groups, n.s. = not significant, * = p<0.05, ** = p<0.01.

Tissue dwelling stages are immobilised/damaged/destroyed by antibody-bound cells within the granulomas (reviewed in ^11^). To quantify antibody levels within granulomas, we isolated worm containing (acute) and non-worm containing (chronic) granulomas and measured antibody levels on the isolated cells by flow cytometry (Sup Fig 2 for gating strategy). Acute granulomas from trickle infected animals had the highest levels of CD11b^+^IgG_1_ ^+^ cells (Fig 4B and 4C). We confirmed these findings by staining for antibodies in intestinal tissue sections isolated from infected mice (Fig 4D-F). Interestingly, we only detected IgG_1_ within the acute granulomas, and did not observe any IgG_2c_ (Fig 2E), which has previously been linked to trapping developing worms within granulomas upon secondary infection ^15^. We did not observe IgA or IgE (Fig 2F) within the granulomas either, confirming that IgG_1_ is the key antibody subtype associated with resistance in our model. Our results are in accordance with previous work studying vaccination mediated protection ^14^.

Chronic granulomas had very low/background levels of IgG_1_ (Fig 4G-I). However, despite this, the granulomas from trickle infected animals were bigger (Fig 4J), with necrotic centres of similar size (Fig 4K). Since this difference was not apparent at D28, the observation is likely due to granuloma age. Once the worms have left the granulomas and/or been killed within it, granulomas resorb. Chronic granulomas in the bolus infected animals are all ∼18 days old (granulomas start forming within 3 days of infection), those in the trickle infected animals range in age from 8-18 days.

To assess whether the differences in granuloma antibody levels were due to differences in antibody production, we measured plasma cell antibody levels at D21 in the MLN and PP, the closest antibody-producing sites to the granulomas (Fig 5, Sup Fig 3 for gating strategy). Plasma cell percentages in the MLN and PP of trickle infected mice were decreased compared to bolus-infected mice; their numbers in the MLN followed this trend (Fig 5A-D). We focused on IgA and IgG_1_ levels because previous work has demonstrated that, in a secondary infection, animals lacking these antibodies have higher worm burden than wildtype animals ^18^. At D21, intracellular IgG_1_ ^+^ plasma cell percentages in the MLN (Fig 5E) and PP (Fig 5F) in trickle infected mice were increased compared to naïve animals but decreased compared to bolus infected animals. Cell numbers in the MLN followed this trend (Fig 5G). As expected, percentages in the MLN were far greater than in the PP ^18^. The decrease could be attributed to a lack of parasite stimulation in the trickle infected animals, which have significantly lower worm burdens. No differences were found between trickle and bolus infected animals with regards to IgA^+^ plasma cell percentages and/or numbers in the MLN and/or PP (Fig 5I-K). The amount of soluble antibody produced per cell, measured by mean fluorescence index (MFI) ratios, did not differ for either the antibody type (IgG_1_ or IgA) or the immune site (MLN or PP) between bolus and trickle infected mice (Fig 5L-O).

**Figure 5:**
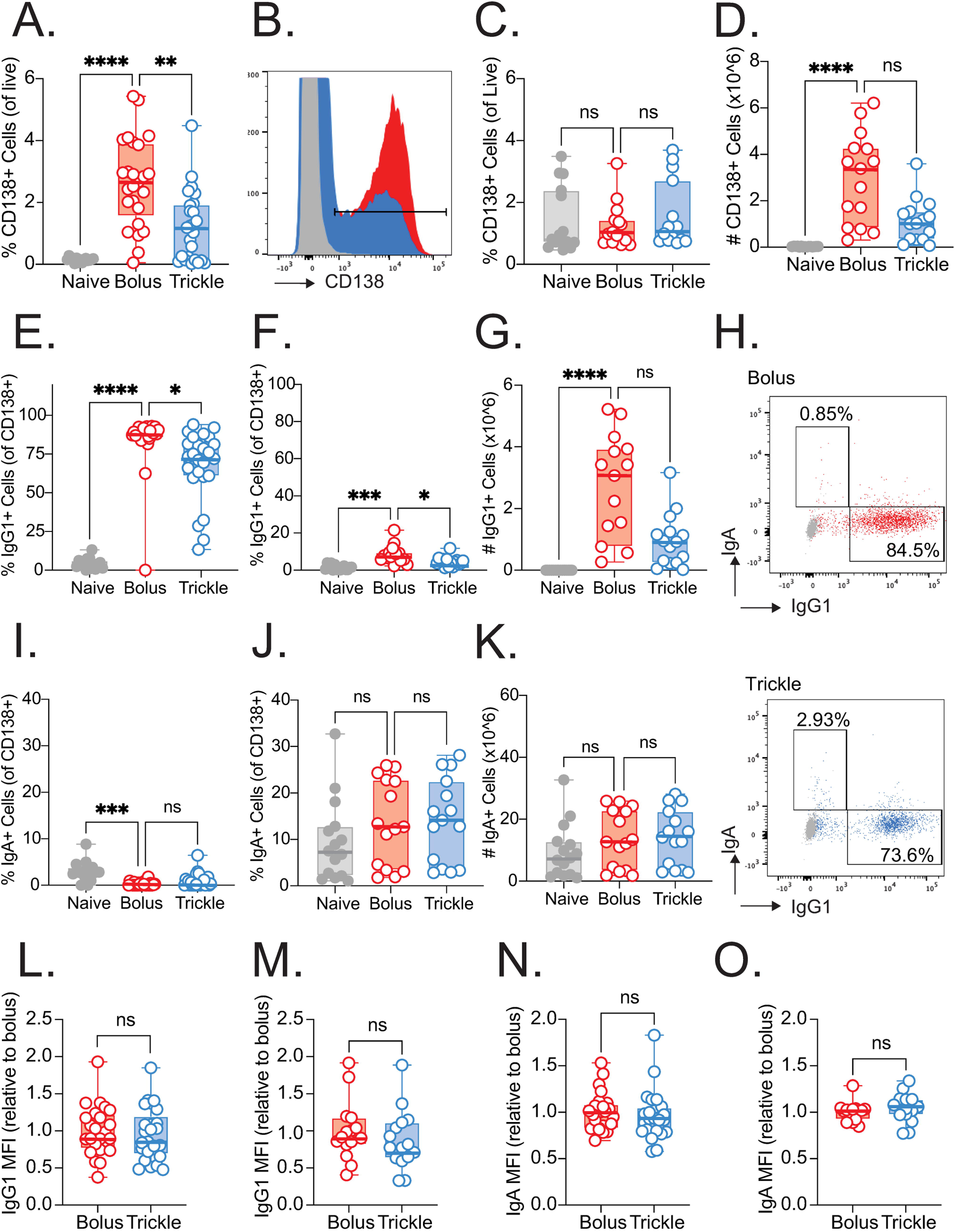
MLN and PP plasma cell antibody production during H. bakeri infection in C57Bl/6 mice. C57Bl/6 mice were infected with 200 *H. bakeri* larvae according to the bolus and trickle infection regimes. Mice were euthanized at D21. Mesenteric lymph node (MLN) and Peyer’s Patches (PP) were isolated to obtain single cell suspensions which were analysed by flow cytometry. (A) The proportion of MLN CD138^+^ (plasma) cells. (B) Representative histogram showing CD138 expression. (C) The proportion of PP CD138^+^ cells. (D) The number of MLN CD138^+^ cells. The proportion of MLN (E) and PP (F) CD138^+^ cells expressing intracellular IgG_1_. (G) The number of MLN CD138^+^ cells expressing intracellular IgG1. (H) Representative flow plots showing IgA and IgG expression on MLN CD138^+^ cells. The proportion of MLN (I) and PP (J) CD138^+^ cells expressing intracellular IgA. (K) The number of MLN CD138^+^ cells expressing intracellular IgA. The IgG_1_ MFI of MLN (L) and PP (M) CD138^+^ cells expressing intracellular IgG1. The IgG MFI of MLN (N) and PP (O) CD138^+^ cells expressing intracellular IgG1. Graphs represent pooled data from a minimum of 3 experiments, bars represent the median, with a minimum of 5 mice per group per experiment. A normality test (D’Agostino & Pearson) was performed, followed by a Kruskal Wallis/Anova (with post tests) and Mann Whitney/T-test to test for statistical significance between trickle and bolus groups, n.s. = not significant, * = p<0.05, ** = p<0.01, *** = p<0.001, **** = p<0.001.

We have already demonstrated that IgG_1_ serum and granuloma levels and serum IgA are undetectable in bolus infected animals at the tissue worm stage (D7), when acute granulomas are present ^13^. This contrasts with trickle infected animals, which have acquired partial resistance (Fig 1), have elevated levels of IgG_1_ ^+^ PP and MLN plasma cells (Fig 5), and increased levels of serum and granuloma IgG_1_ (Fig 3 and 4) during tissue worm development. However, when we passively transferred serum from trickle infected mice to bolus infected animals, we did not detect IgG_1_ within their acute granulomas (Sup Fig 4), suggesting that cells, and not only antibodies, present in the granuloma have an important role to play. This supports the idea that sterile immunity may require either extremely high antibody titers, or the activation of additional protective effector cell populations ^14^.

IgG-bound macrophages and eosinophils are the cells of the granuloma responsible for parasite damage/death (reviewed in ^11^). We found that within the IgG_1_ ^+^ cells of the trickle infected animals, SiglecF^+^ (eosinophils) and CD206^+^ (macrophages) cells accounted for approximately 50% of the antibody bound cells (Fig 6A, Sup Fig 2 for gating strategy). When comparing the composition of the whole acute granulomas from trickle and bolus infected animals, trickle infected animals had a far higher proportion of SiglecF^+^ (Fig 6B) and CD206^+^ cells (Fig 6C) than bolus infected mice. Proportions of Ly6G^+^ (neutrophils, Fig 6C) and NK1.1^+^ (NK cells, Fig 6D) cells remained unchanged between the two groups.

**Figure 6:**
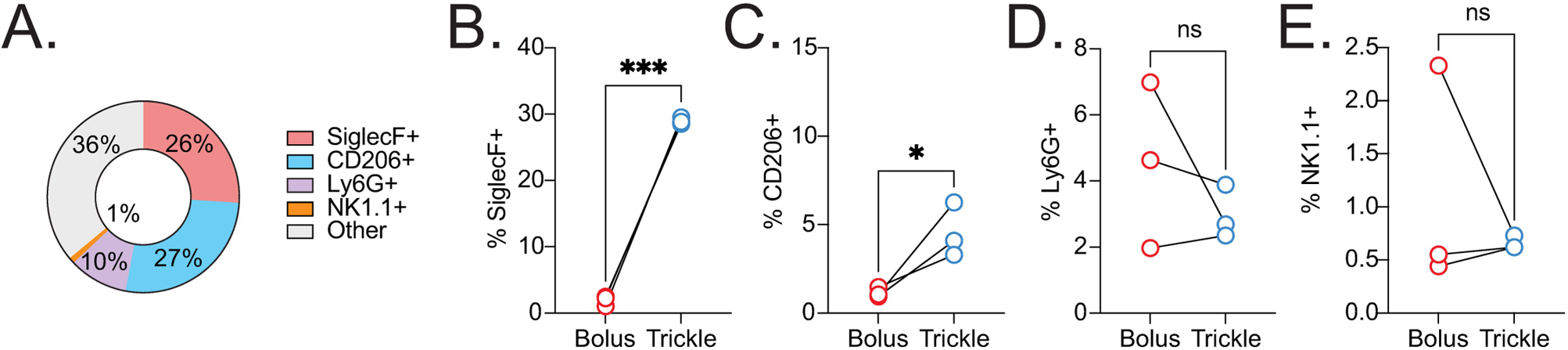
Acute granuloma cell composition. C57Bl/6 mice were infected with 200 *H. bakeri* according to bolus and/or trickle models of infection and euthanised at D21. Granulomas were isolated from the small intestine of 5 mice per group. Granulomas from the individual animals were pooled to form one sample per group. Cells were obtained by tissue digestion of the pooled samples and surface stained by flow cytometry. (A) Proportions of IgG -bound cells (SiglecF^+^, CD206^+^, Ly6G^+^ and NK1.1^+^) were calculated within the acute (worm containing) granulomas of trickle infected mice. Numbers reflect the mean of five independent experiments. Proportions of acute granuloma (B) SiglecF^+^, (C) CD206^+^, (D) Ly6G^+^ and (E) NK1.1^+^ cells within bolus and trickle infected mice were calculated for three independent experiments. A paired T-test to test for statistical significance between trickle and bolus groups was performed, n.s. = not significant, * = p<0.05, *** = p<0.001.

Resistance in trickle infected mice is therefore associated with the presence of both antibodies and SiglecF^+^/CD206^+^ cells at the host/parasite interface during tissue development. However, if repeated infections provide protection from Hb, why is this not reflected in nature, where repeated infections are common? Wild rodents have a high incidence of Hb ^25,26^ and, more generally, intestinal worms are common parasites of humans ^1^, livestock (reviewed in ^2^) and pets ^3^.

### Resistance to Hb in trickle infected animals is lost when they are pre-infected with Tg

In natural settings, co-infection is common ^28^. This translates to complex host immune environments. To integrate the increased complexity produced by a mixed immune response and potential co-infections into our research, we used *Toxoplasma gondii* (Tg) as a potent inflammatory parasite. The order and time between infections was optimized to ensure minimal pathology while retaining characteristic immune signatures: animals were pre-infected with 10 Tg cysts 14 days prior to trickle Hb infection (Fig 7A). In a previous study using the bolus model of infection, pre-infection with Tg skewed the immune response to a mixed Th1/Th2 Hp immune response by D28. Focusing on the sections with the most granulomas (first half of the small intestine, with SI1 equating to the duodenum and SI2 to the first part of the jejunum), at D21, we found that intestinal levels of *ifnγ* and *il10* gene expression were increased in the jejunum of HbTg trickle infected mice, while *il13* levels were not different (Fig 7B-G). We wanted to determine whether this more complex immune environment would disrupt the effective anti-larval immunity observed during trickle infection, and impact worm burden.

**Figure 7:**
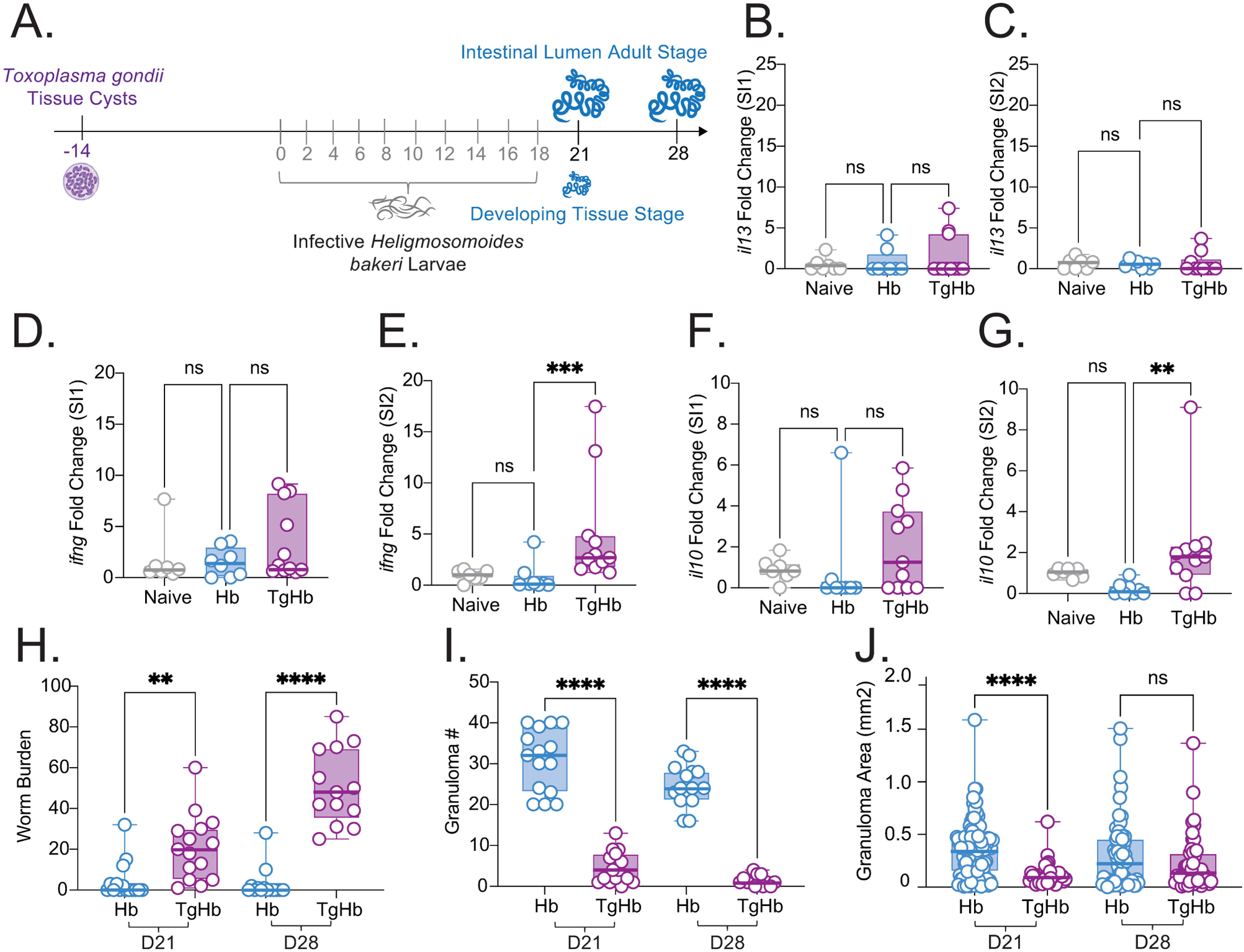
The T. gondii/H. bakeri trickle coinfection model in C57Bl/6 mice. C57Bl/6 mice were infected with 200 *H. polygyrus* according to the coinfection and trickle infection regimes. (A) Coinfection trickle infection regime: mice are infected with 20 Me49 *T. gondii* cysts 14 days prior to *H. bakeri* infection which consists of 200 larvae in total, but in multiple doses over the course of infection (in grey). Doses of 20 larvae are trickled on days 0, 2, 4, 6, 8, 10, 12, 14, 16 and 18 post-infection Animals are euthanised on D21 (3 day window after the final dose), where tissue and lumen parasites can be observed. Animals are also euthanised on D28 (10-day window after the final dose), where only lumen parasites are found. Mice were infected according to the Hb (blue) and TgHb (purple) regimes. (B-G) RNA was extracted was from the small intestine, which had been cut into four equal portions. The duodenum (SI1) and jejunum (SI2) were used. Gene expression was measured by quantitative RT-PCR for *il13* (B & C), *ifnγ* (D & E) and *Il10* (F & G). Adult worms (H) and granulomas (I) were counted in the small intestine at D21 and D28. (J) Granuloma area in C57Bl/6 mice was measured on H and E stained sections from whole small intestine swiss rolls. Graphs represent pooled data from a minimum of 2 experiments, bars represent the median, with a minimum of 4 mice per group per experiment. A normality test was performed, followed by a Kruskal Wallis/Anova (with post-tests) and Mann Whitney to test for statistical significance between Hb and HbTg groups, n.s. = not significant, ** = p<0.01, ***= p<0.001, **** = p<0.0001.

In this context, we found that pre infection with Tg led to higher worm burdens at both D21 and D28 (Fig 7H). Unsurprisingly, the increased parasite burden was associated with fewer granulomas (Fig 7I) of smaller size (Fig 7J). Taking a closer look at the tissue granulomas from TgHb animals, we found they had decreased levels of IgG_1_ ^+^ cells at D21, as observed by flow cytometry (Fig 8A and 8B) and immunofluorescence (Fig 8C). The overall cellular composition of granulomas was also altered: proportions of SiglecF^+^ and CD206^+^ cells were decreased (Fig 8D and 8E); proportions of Ly6G^+^ (Fig 8F) and NK1.1^+^ (Fig 8G) cells remained unchanged. Histology confirmed our flow cytometry findings: HbTg animals had fewer eosinophils (Fig 8H). Interestingly, total macrophages numbers were similar between the two groups (Fig 8I). However, this is likely because similar numbers of macrophages are recruited to the granulomas in both groups. However, in HbTg animals, the influx of CD206+ macrophages is reduced is counterbalanced by the influx of other more inflammatory macrophages.

**Figure 8:**
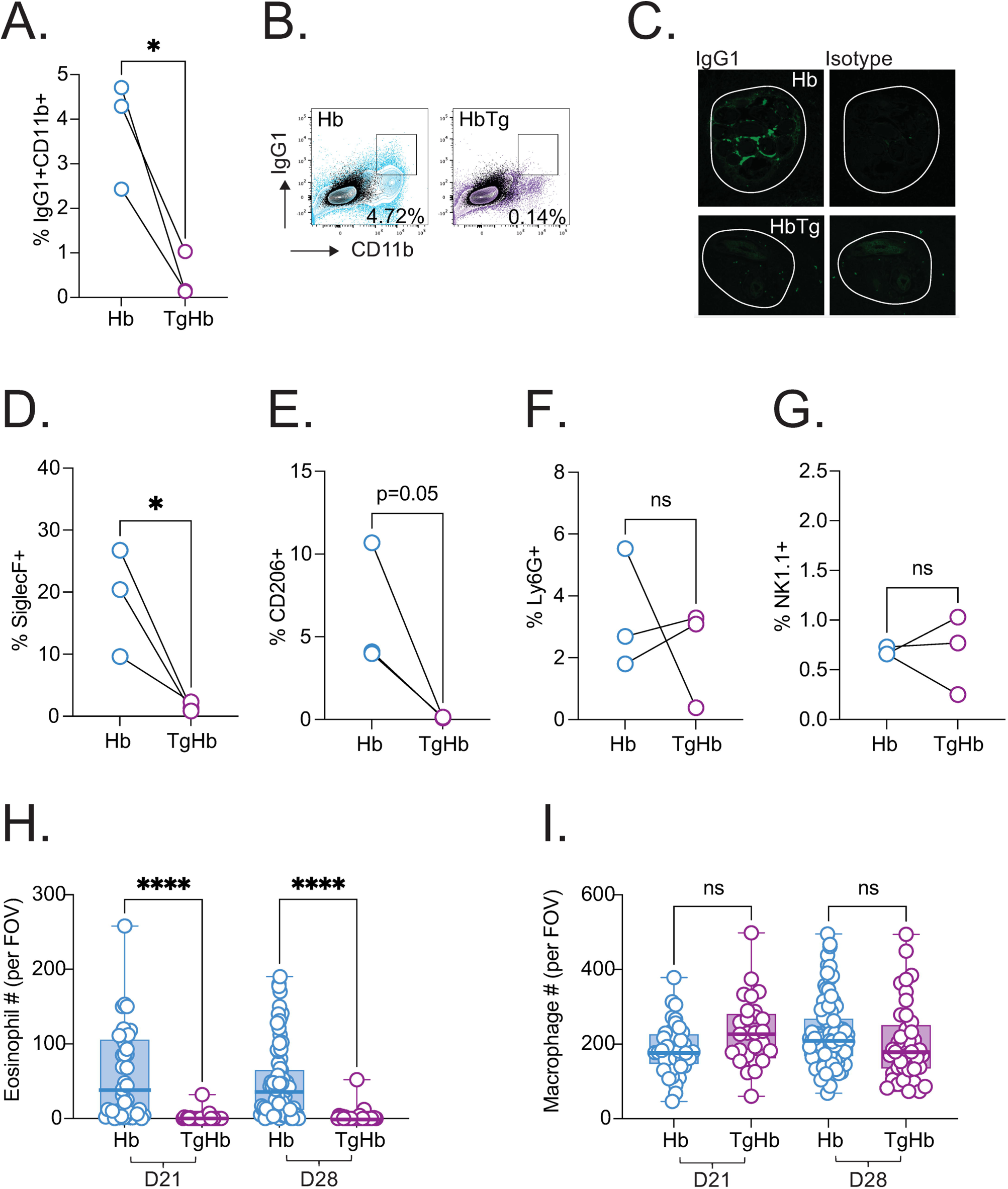
Granuloma characteristics in the T. gondii/H. bakeri trickle coinfection model in C57Bl/6 mice. C57Bl/6 mice were infected with 200 *H. bakeri* according to the trickle single or co-infection models, and euthanised on D21. Co-infected animals are infected with 10 cysts of Me49 *T. gondii* 14 days before the first *H. bakeri* dose. Granulomas were isolated from the small intestine of 5 mice per group. Granulomas from the individual animals were pooled to form one sample per group. Cells were obtained by tissue digestion of the pooled samples and stained by flow cytometry. (A) The percentage of CD11b^+^IgG ^+^ cells as a proportion of live granuloma cells (with T and B cells removed) from acute granulomas was calculated for three independent experiments. (B) Representative flow cytometry plots showing IgG_1_ and CD11b expression on live granuloma cells from acute granulomas. (C) Formalin fixed, paraffin-embedded sections were obtained from small intestine swiss rolls. These sections were co-stained with DAPI and anti-mouse IgG_1_. Scale bar=100μm. Antibody stain (left), isotype control stain (right). Representative granulomas (white lines) containing worms (acute granulomas) from trickle and co-infected mice. (D) Proportions of acute granuloma (D) SiglecF^+^, (E) CD206^+^, (F) Ly6G^+^ and (G) NK1.1^+^ cells within trickle and co-infected mice were calculated for three independent experiments. A paired T-test to test for statistical significance between two groups was performed, n.s. = not significant, * = p<0.05. Granulomas were identified on H and E stained slides of whole small intestine. Eosinophils (H) and macrophages (I) were counted within the center of the granulomas at x400 magnification. Cells were counted for one field of view (FOV) per granuloma. A normality test was performed, followed by a Kruskal Wallis/Anova (with post-tests) to test for statistical significance between trickle and co-infected groups, n.s. = not significant, * = p<0.05, **** = p<0.001.

To study whether reduced antibody production was responsible for reduced granuloma antibody levels, we measured plasma cell antibody levels at D21 in the MLN using flow cytometry (Fig 9). While total MLN cell numbers were decreased in co-infected animals (Fig 9A), plasma cell percentage and numbers were increased (Fig 9B and 9C). The proportion of MLN IgG_1_ ^+^ plasma cells was also increased in TgHb animals (Fig 9D), although their number and antibody production level per cell (MFI) were similar between the two groups (Fig 9E and 9F). However, when focusing on antigen specific IgG_1_ producing cells using ELIspot, TgHb animals had far fewer cells than Hb animals: a maximum of 93 vs a mean of 318 (Fig 9G). The difference was not apparent when measuring total IgG_1_ (Fig 9H). Finally MLN IgA^+^ plasma cells were decreased in number (Fig 9I) but not in proportion (Fig 9J) or in their capacity to make IgA (MFI, Fig 9K). Therefore, in co-infected animals, increased worm burdens are associated with a reduction in the efficacy of granulomas: they lack the presence of SiglecF^+^ and CD206^+^ cells as well as parasite specific IgG_1_.

**Figure 9:**
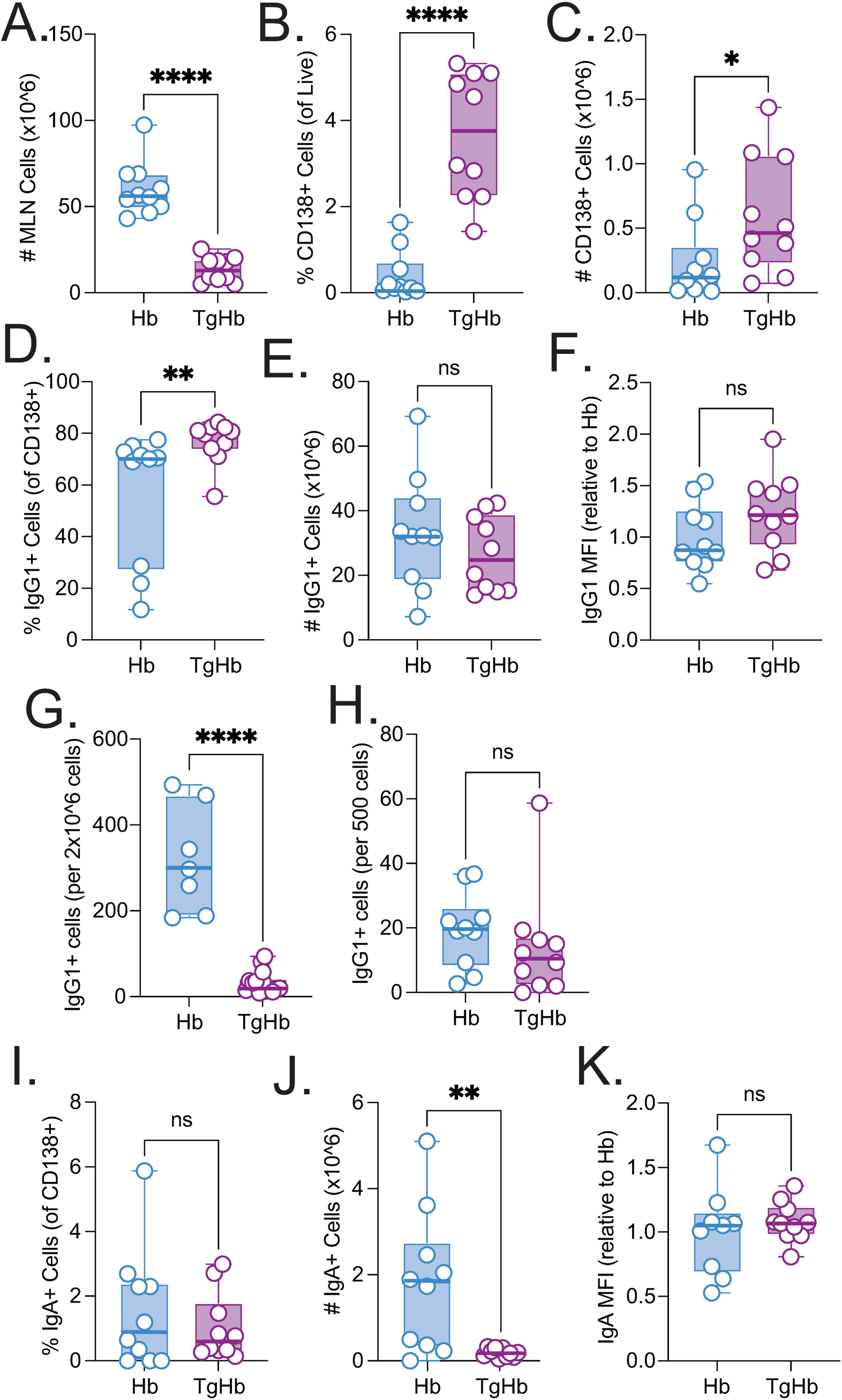
Plasma cell and antibody dynamics in T. gondii/H. bakeri trickle co-infected C57Bl/6 mice. C57Bl/6 mice were infected with 200 *H. bakeri* according to the trickle single or co-infection models, and euthanised on D21. Co-infected animals are infected with 10 cysts of Me49 *T. gondii* 14 days before the first *H. bakeri* dose. Mesenteric lymph nodes (MLN) were isolated to obtain single cell suspensions which were analysed by flow cytometry. (A) The number of total viable MLN cells. (B) The proportion of CD138^+^ (plasma) cells. (C) The number of CD138^+^ cells. (D) The proportion of CD138^+^ cells expressing intracellular IgG. (E) The number of CD138^+^ cells expressing intracellular IgG. (F) The IgG MFI of CD138^+^ cells expressing intracellular IgG. The number of MLN *H. bakeri* -specific (G) and total (H) IgG_1_ producing cells assessed by ELISPOT. (I) The proportion of CD138^+^ cells expressing intracellular IgA. (J) The number of CD138^+^ cells expressing intracellular IgA. (K) The IgA MFI of CD138^+^ cells expressing intracellular IgG_1_. Graphs represent pooled data from a minimum of 2 experiments, bars represent the median, with a minimum of 5 mice per group per experiment. A normality test (D’Agostino & Pearson) was performed, followed by a Kruskal Wallis/Anova (with post-tests) and Mann Whitney/T-test to test for statistical significance between Hb and TgHb groups, n.s. = not significant, * = p<0.05, ** = p<0.01, **** = p<0.001.

## DISCUSSION

The pattern of resilience/resistance observed in the laboratory is not what is observed in naturally infected populations of animals – instead, GINs are widespread in wild populations. As an example, studies in wild mice have reported a greater than 60% prevalence with one or more helminth parasites ^25–27^ and in a recent Albertan survey, 100% of cattle herds were found to be infected with intestinal nematodes ^40^. This is likely due to stark differences in laboratory conditions compared to natural environments. Factors such as nutrition, age, genetics and environment all play a role. Importantly, so do immune history and co-infection, which can affect the immune system’s ability to respond to a parasite infection.

GIN infections typically occur over extended periods of time, with small amounts of parasite taken up regularly during feeding. The mode of infection is an important factor in the ensuing immune responses. Multiple low dose, or trickle infections, have long been associated with increased resistance to helminth parasites, resulting in a “resistant”/resilient phenotype in animals that are normally considered susceptible ^13,41–44^. This acquired resistance can remain long-term, with trickle infected animals maintaining their ability to eliminate worms during subsequent challenge infections ^41^. Here, we show that effective granulomas are key to eliminating incoming GIN larvae: anti-GIN antibodies must be produced in time to target tissue developing parasites and directed to the host/parasite interface where they can bind activated myeloid cells to immobilise/damage/kill the parasites. Bolus and TgHb trickle infected animals had high worm burdens. In bolus-infected susceptible animals, the antibodies were not produced in time to reach the acute granulomas before the worm exited the tissue. In TgHb trickle-infected animals, granuloma formation was impacted by a reduction in the influx of GIN-specific antibody and myeloid cells to the host-parasite interface.

Granulomas are formed as a response to tissue invading Hb and consist mainly of macrophages and eosinophils. The exact mechanisms by which tissue developing parasites are immobilised/damaged/destroyed within the granulomas *in vivo* have not been fully characterised. A study involving *Nippostrongylus brasiliensis* (closely related to Hb) infection in mice found that following primary inoculation with the parasite, Th2-primed macrophages remained in the lung (site of parasite development) and facilitated rapid larval clearance upon reinfection ^45^. Trickle infection of *N. brasiliensis* in its natural host, the rat, results in increased lung macrophages and eosinophils ^46^. We found increased levels of CD206^+^ cells in the tissue granulomas of trickle infected mice suggesting that long-lived, previously sensitized CD206^+^ macrophages may also play a direct role in the clearance of Hb tissue stages from the intestine in these animals. During *Litomosoides sigmodontis* infection, a parasite which does not cause granulomas but develops in the pleural cavity, CD206^+^ pleural cavity macrophage populations increase due to the proliferation of tissue-resident macrophages and, to a lesser extent, the re-programming of recruited inflammatory macrophages ^47^. One caveat of our work is that we did not assess markers of tissue residence or recruitment on granuloma macrophages. Without this information, no definite conclusion can be made about the origins of the cells that are required for the effective anti-larval immune response seen in trickle-infected animals.

*In vitro* and in the presence of immune serum, CD11b on bone marrow derived macrophages can directly bind Hb larvae using complement 3 ^16^. This allows FcγR1 to interact with parasite bound IgG (but not IgG_1_) antibodies to immobilize the larvae ^15^. However, while immune serum allows macrophages to bind to Hp larvae *in vitro*, it does not impact infectivity, suggesting that macrophages are important only in parasite immobilisation and not damage/killing. The authors hypothesised that this rapid activation via IgG_2a/c_ and Fc receptors would likely be important against tissue dwelling parasites during acute infection, but this has yet to be proved as the main mechanism of parasite immobilisation *in vivo*. In accordance with the authors, we observed CD11b^+^ antibody-bound macrophages within the worm-containing granulomas of trickle-infected mice. However, only IgG_1_, and not IgG2_c_, was observed in the granulomas which supports previous reports identifying IgG_1_ as the protective mechanism in vaccinated mice ^48^. Despite their proximity to Hb’s infective niche, we saw no increase in IgG ^+^ plasma cells in the PP of Hb infected animals. This is likely due to parasite-derived immunomodulatory molecules preventing Th2 polarization at this site, as PP in direct contact with Hb larvae show significant expansion of Treg populations ^49^.

A recent systematic review has concluded that IgA is associated with partial protection from helminths in mice, rats and sheep ^50^. Mechanism of protection include entrapment, neutralization, and eosinophil-mediated ADCC. Mice infected with Hb have elevated IgA levels during primary ^17^ and repeated ^44^ infections; mice lacking IgA and infected with Hb in a drug cure/reinfection model do not clear parasites as effectively as wildtype animals ^18^. We did not observe IgA within the granulomas of trickle infected animals and did not associate IgA with tissue parasite immobilisation/damage/ clearance.

The role of eosinophils during helminth infection has long been debated (reviewed in ^51^). *In vitro* studies have demonstrated that peritoneal and bone marrow derived eosinophils can respectively adhere to ^52^ and limit the motility of parasitic worm larvae through extracellular traps ^53^. Eosinophils obtained from the mammary washes of sheep can kill *Haemonchus contortus* (closely related to Hb) *in vitro* in the presence of antibody; efficacy was increased in the presence of complement and by using ‘primed’ eosinophils from previously infected sheep ^54^. *In vivo,* studies using transgenic animals show that killing by toxic granules is also important: EPX- and MBP1-deficient Balb/c mice have higher worm burdens than wildtype mice after *L. sigmodontis* infection ^55^. Data from *H. contortus* immunised sheep supports these findings: tissue larvae in the vicinity of eosinophils were found to be at varying stages of damage ^56^. However, in Hb infected mice, parasite loads in mice lacking eosinophils (Balb/c Δdbl-GATA-1 and C57Bl/6 PHIL) are similar to wild type animals ^57^ and others have shown that eosinophils do not adhere to Hb *in vitro* ^16^, questioning the importance of this cell type in parasite killing. Despite having similar worm burdens, Δdbl-GATA-1 mice challenged with Hb after vaccination had significantly higher numbers of live larvae trapped in their intestinal wall compared to their wild-type counterparts ^48^. This suggests that eosinophils are responsible for killing but not immobilising tissue-dwelling parasite stages. The cells likely work in concert with macrophages to limit the number of tissue dwelling parasites that develop into adults, which explains their elevated numbers in the granulomas of trickle infected animals.

We have previously shown that by 10 days of infection with Tg, a strong increase in IFNγ and IL-10 develops along the whole small intestine, the PP, MLN, SPL and the liver ^58^. This immune environment impacts the ability of C57Bl/6 mice to mount an effective immune response against a trickle Hb infection. We observed fewer and smaller granulomas; they contained a lesser proportion of CD206^+^, SiglecF^+^ and IgG -bound cells. To the naked eye, the acute granulomas were not as well formed, and the chronic granulomas did not have the characteristic bead-like appearance that is associated with bolus/trickle infections. Interestingly, in wild mice, granulomas are rarely observed by eye (Pers. Comms. Dr Simon Babayan). This could be due to differences in parasite lifecycles (*H. polygyrus* vs. *H. bakeri*), host responses (wood mice vs. laboratory mice) or could suggest that a more complex immune environment impacts the formation of these structures.

In a model of allergic inflammation, pre-infection with Tg resulted in decreased serum allergen-specific IgG levels ^59^. Despite similar levels of IgG_1_-producing plasma cells and IgG production, HbTg infected animals also had reduced antigen specific (Hb) IgG ^+^ MLN plasma cell levels compared to Hb trickle infected mice, which helps explain why we observed very little IgG_1_ ^+^ in the granulomas of TgHb infected mice. These results reflect gastrointestinal nematode infections in the wild whereby *H. polygyrus* infected mice co-infected with the coccidian *Eimeria* had lower *H. polygyrus*-specific serum IgG ^+^ titers than *H. polygyrus* only infected mice ^26^. HbTg infected animals also had elevated proportions and numbers of plasma cells, indicating the induction of a polyclonal antibody response against both pathogens, rather than a strong Hb-specific response. The production of polyclonal antibodies during Hb infection has previously been shown to reduce fecundity of adult worms in the intestinal lumen but not to reduce worm burden ^18^, supporting our findings.

Despite receiving the same multiple small dose infection as trickle infected animals, the proportion of SiglecF^+^ and CD206^+^ cells in the granulomas from co-infected animals was severely reduced, indicating that Tg significantly impaired the recruitment of these cells to the granulomas. TgHb bolus infected mice have been shown to simultaneously decrease proportions of Th2 (GATA3^+^CD4^+^) cells and drastically increase proportions of Th1 (T-Bet^+^CD4^+^ and T-Bet^+^CD8^+^) cells in the small intestine, mesenteric lymph nodes, and spleen ^30^. Hb-specific CD4^+^ T cells produce IFNγ rather than IL-4 or IL-13 *in vitro.* This observed shift would also impact other cells. For example, intestinal epithelial cells recruit eosinophils to the small intestine through the production of eotaxin-1, following activation in a Th2 environment ^60^. The impairment of Th2 cytokine production would therefore directly affect the recruitment and activation of eosinophils and CD206^+^ macrophages, both of which are dependent on Th2 cytokine signaling (reviewed in ^61,62^). This has also been observed in the context of *N. brasiliensis*. When pre-infected with the apicomplexan *Plasmodium chabaudi* (related to Tg), *N. brasiliensis* co-infected animals had reduced CD206^+^ macrophage accumulation at the site of parasite development, the lung ^63^. When Tg was the pre-infecting parasite, increased shedding of eggs (both in amount of eggs and length of time) was observed in the co-infected animals ^64^.

Increased worm fecundity was also observed in TgHb infected animals using the bolus infection model ^30^, but unlike with the trickle model, no difference in worm burden was detected.

Recent work has demonstrated that larval trapping after secondary infection does not require macrophages, rather it is intestinal epithelial cells that play a critical role in this regard ^65^. The exact molecules involved remain to be identified, although the process was reliant on IL-4R signalling. The TgHb mice had significantly increased levels of intestinal *ifnγ* levels throughout the experiment which may have hampered the IEC IL-4R signalling required for effective clearance.

NK and neutrophils proportions did not change with the mode of infection. Recent work has argued that NK cells are essential for host protection during the acute phase of Hb infection. Removing NK cells did not impact worm burden or fitness in the bolus mode of infection, rather intestinal injury was increased ^66^. Ly6G^+^ cells have been identified within the granulomas of Hb infected animals ^67,68^. However, immunity to Hb following vaccination was unaffected by the loss of neutrophils induced by antibody treatment in C57Bl/6 animals ^48^, and neutrophil populations did not increase in the lungs of mice with a secondary *N. brasiliensis* infection ^45^.

One limitation of our study is that we did not assess whether differences in the microbiome could account for the changes we observed. In the laboratory setting, Tg infection (Me49, oral gavage) leads to dysbiosis, characterised by an increase in proinflammatory Pseudomonadota species and a reduction of beneficial commensal species, that persists into the chronic stage of infection ^69^. By contrast, Hb infection in C57Bl/6 mice leads to duodenal increases in the abundance of the immunoregulatory *Lactobacillus* species ^70^ and infections in germ free mice result in increased worm burdens ^71^. However, in experiments using the trickle, rather than bolus, infection model it was demonstrated that mice infected with *Trichuris muris* recovered alpha diversity following the development of resistance ^41^. Interestingly, in mice infected with *T. muris* which were then moved outdoors and acquired an enriched microbiome, the authors measured a decrease in Th2 and increase in Th1 immunity associated with increased worm burdens and biomass ^72^, somewhat reflecting the more nuanced immune environments created in our co-infection model. However, it is also important to remember that other factors, unrelated to host biology, such as temperature and humidity which are strong moderators of the microbiome ^73^, may play a significant role in infection outcome in the wild. And that restricting studies to Th1 vs. Th2 in a particular setting at a particular timepoint likely oversimplifies the mechanisms underlying the optimal host response ^74^. However, compared to both the typical experimental infection (bolus) and the single infection grazing type infection (trickle), the co-infection trickle model described here more realistically reflects immune responses in real-world infections. Feeding through grazing exposes hosts to a variety of pathogens. While our two-parasite system is not nearly as complex as real-world infections, our research begins to explain the overwhelming prevalence of helminth infections worldwide.

## ACKNOWLEDGEMENTS

This works was funded through Dr Finney’s grants from the Canadian Foundation for Innovation and the Natural Sciences and Engineering Research Council of Canada (NSERC), as well as scholarships for Drs Anupama Ariyaratne (NSERC Create in Host Parasite Interactions), Shashini Perera (Alberta Graduate Excellence Scholarship), Naomi Chege (Graeme Bell and Norma Kay Sullivan-Bell Graduate Scholarship in Biology), Emma Forrester (University of Calgary PURE Scholarship), Mayara Luzzi (Mitacs Globalink), Joel Bowron (NSERC), Edina Szabo (UCalgary Eyes High). The funders had no role in study design, analysis or reporting.

## DATA ACCESS

All data not available in the manuscript or supplemental materials are available from the corresponding author upon request.

---

***Supplemental Figure 1: Characteristics of the H. bakeri trickle infection model in Balb/c mice.*** Balb/c mice were infected with 200 *H. bakeri* according to the bolus and trickle infection regimes. Adult worms (A) and granulomas (B) were counted in the small intestine at D28. Mice were infected according to the bolus (red) and trickle (blue) regimes. Single cell suspensions were isolated from the spleen (SPL) and mesenteric lymph nodes (MLN) and cultured for 48 hours in the presence of Hb antigen. MLN/SPL cell numbers were calculated (C & F), and IL-4 (D & G) and IL-13 (E & H) cytokine levels were measured in the supernatant by ELISA. (I) Total serum IgE, (J) total serum IgG_1_ and (K) parasite-specific serum IgG_1_ were measured by ELISA. Levels in naïve controls were undetectable for parasite specific IgG_1_. A normality test (D’Agostino & Pearson) was performed, followed by a Kruskal Wallis/Anova (with post-tests) and Mann Whitney/T-test to test for statistical significance between trickle and bolus groups, n.s. = not significant, ** = p<0.01, *** = p<0.001, **** = p<0.001.

***Supplementary Figure 2: Gating strategy for IgG_1_ ^+^ granuloma cells.*** Granulomas (acute and chronic) were isolated from the small intestine of 5 mice per group. Granulomas from the individual animals were pooled to form one sample per group. Cells were obtained by tissue digestion of the pooled samples and surface stained by flow cytometry. Gating strategy involved removing doublets using FSC and SSC. Dead cells (positive for viability stain) were excluded. B and T cells were excluded by removing CD3^+^ and CD19^+^ cells from the analysis. Antibody-bound cells were identified as CD11b^+^IgG_1_^+^ cells using appropriate controls. To identify specific cell types, eosinophils were identified as SiglecF^+^, macrophages as CD206^+^, neutrophils as Ly6G^+^ and NK cells as NK1.1^+^, using appropriate controls.

***Supplementary Figure 3: Gating strategy for IgG _1_ and IgA producing plasma cells.*** Organs were isolated to obtain single cell suspensions which were analysed by flow cytometry. Gating strategy involved removing doublets using FSC and SSC. Dead cells (positive for viability stain) were excluded. IgD^+^CD3^−^ cells were excluded by removing CD3^+^ and IgD^+^ cells from the analysis. IgG_1_ ^+^CD138^+^ and IgA^+^CD138^+^ populations were identified using appropriate controls.

***Supplementary Figure 4: IgG_1_ ^+^ cells in the granulomas of D7 bolus infected mice.*** Serum was administered intravenously at D0, D2, D4, D6 to bolus infected animals. Formalin fixed paraffin embedded small intestine swiss rolls were obtained on D7 and co-stained with anti-mouse IgG_1_ and DAPI. Top: mice who received trickle immune serum obtained from trickle infected mice. Bottom: mice who received non-immune control serum obtained from naïve animals. Experiment was performed once with 5 mice per group, and representative images are presented. Scale bar=100μm.

