## Supplementary figures and images for "Co-infection with *Toxoplasma gondii* leads to a loss of resistance in *Heligmosomoides bakeri* trickle-infected mice due to ineffective granulomas"

### Supplemental Figure 1

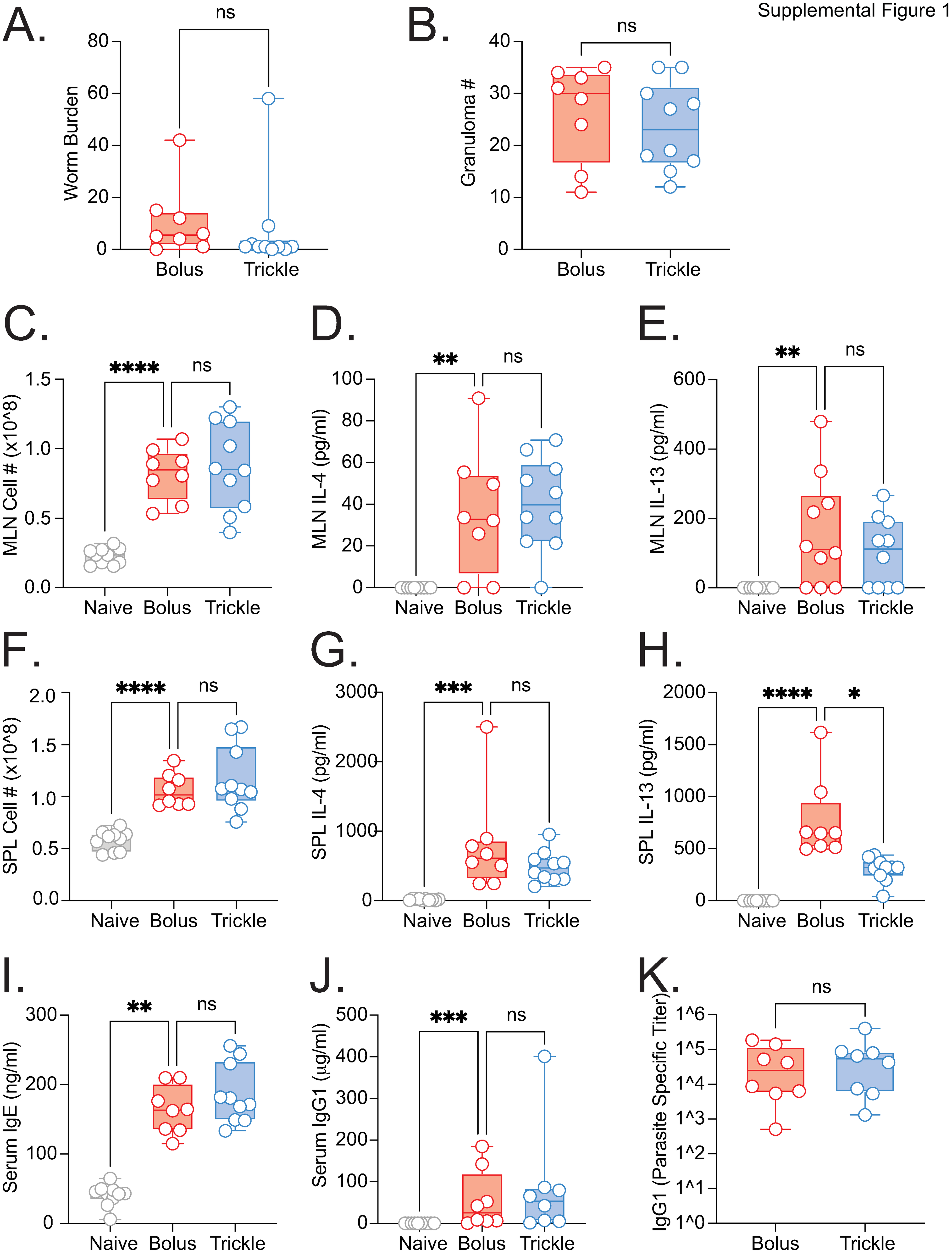

### Supplemental Figure 2

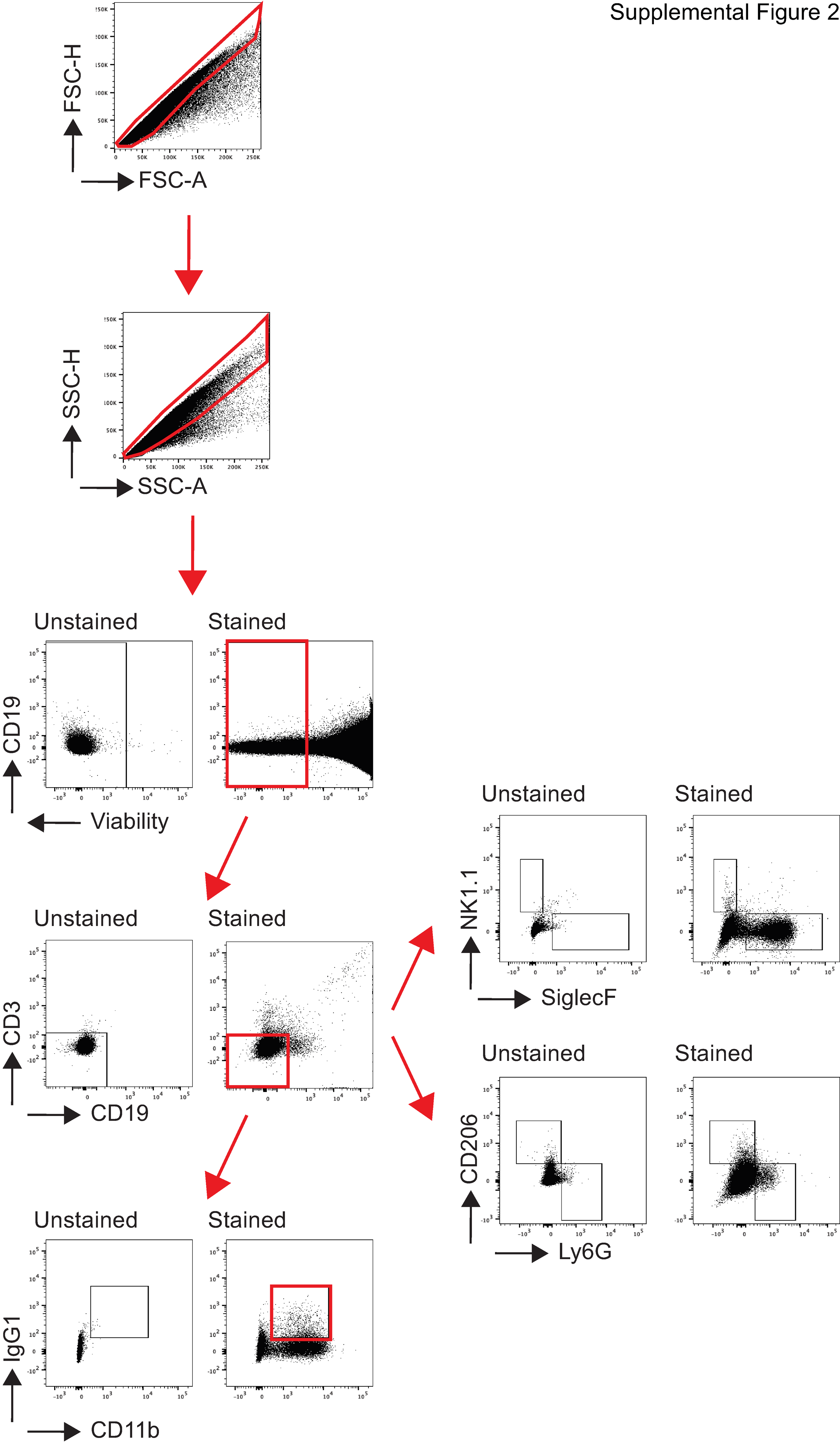

### Supplemental Figure 3

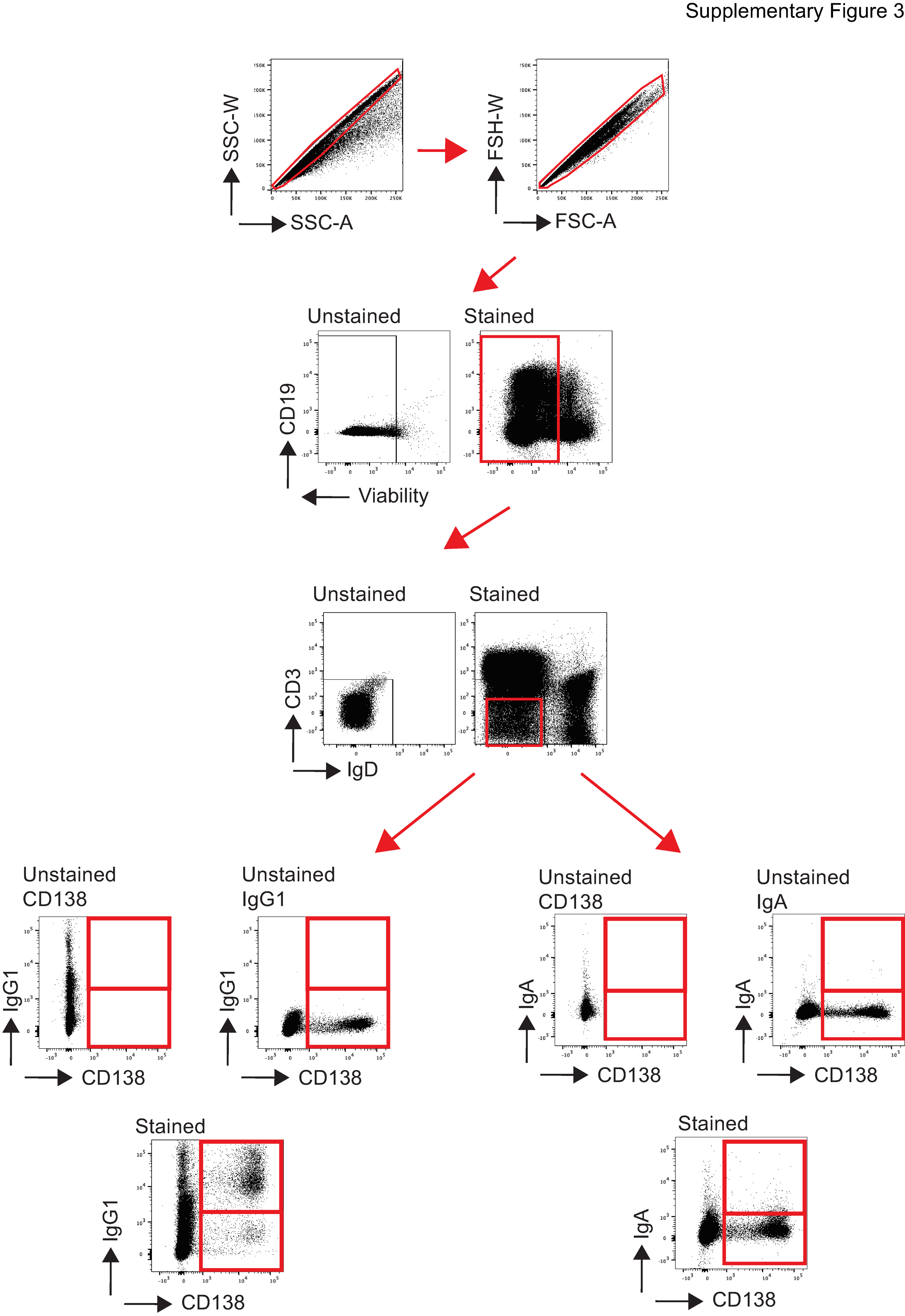

### Supplemental Figure 4

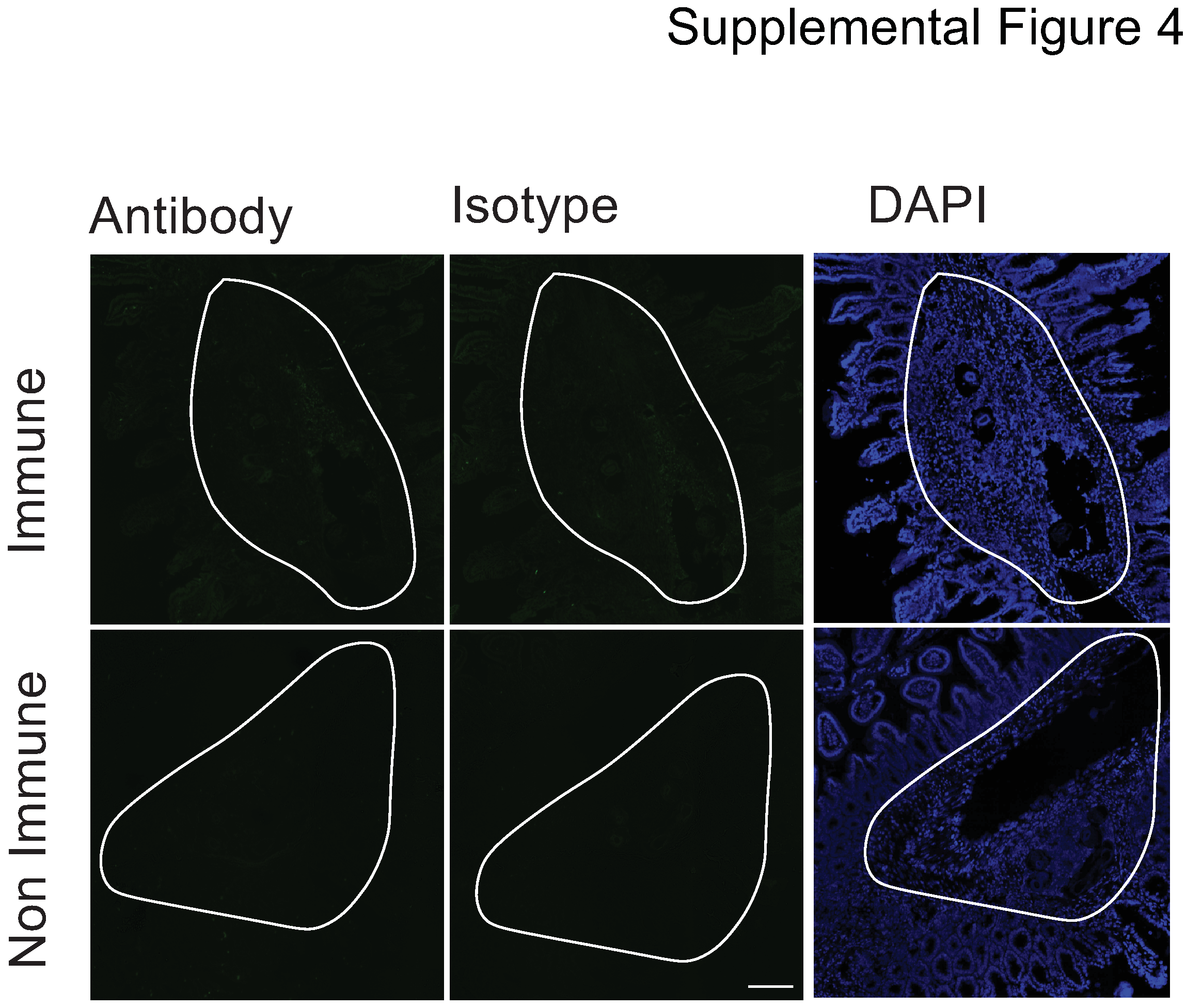
